# Targeting Amygdala-Brainstem Synapses to Reverse Prepulse Inhibition Deficits

**DOI:** 10.1101/2025.09.24.676662

**Authors:** Wanyun Huang, Michelle Kha, Adolfo E. Cuadra, Okoh Frimpong, Laura Snow, Andrea Gouin, Stephanie Padilla, Karine Fénelon

## Abstract

Sensorimotor gating, a fundamental pre-attentive process, can be assessed using the acoustic prepulse inhibition (PPI) assay. PPI deficits are a hallmark endophenotype of schizophrenia and are observed across various neuropsychiatric disorders, often predicting symptoms such as attention impairments, psychosis, and other cognitive/motor dysfunctions. Reversal of PPI deficits is routinely tested in disease models as a preclinical trial for antipsychotic drug screening. However, the cellular and circuit-level mechanisms underlying PPI deficits remain unclear, limiting therapeutic progress. We recently identified an uncharted pathway in mice, by which glutamatergic neurons in the central nucleus of the amygdala (CeA) activate glycinergic neurons in the caudal pontine reticular nucleus (PnC), contributing to PPI regulation. Given the prevalence of amygdala dysfunction in disorders associated with PPI deficits, CeA-PnC glutamatergic synapses represent a novel therapeutic target. Here, using “Cal-Light,” an *in vivo* Ca2^+^-dependent photo-tagging approach, we precisely identified CeA and PnC neurons active during acoustic startle and PPI with high spatiotemporal resolution. Furthermore, we used mice carrying homozygous deficiency in *PRODH*, a schizophrenia-relevant gene encoding proline dehydrogenase and modulating PPI. While *Prodh^−/−^* mice showed aberrant CeA neuronal properties and reduced PPI levels, we restored PPI by photo-activating CeA-PnC glutamatergic synapses, underscoring their involvement in pathological states. These findings provide new mechanistic insights into the amygdala-brainstem circuitry that underlies PPI deficits, offering new potential for therapeutic interventions.

## Introduction

Sensorimotor gating refers to the ability of a sensory event to inhibit or modulate universal behaviors, such as startle responses (Braff and Geyer, 1990; Li et al., 2009; Powell et al., 2012). Sensorimotor gating can be operationally measured through prepulse inhibition (PPI) of the acoustic startle response, with changes in PPI indicating alterations in gating (Graham, 1975; Norris and Blumenthal, 1996; van den Buuse, 2007). PPI is a universal phenomenon that occurs when a weak sensory stimulus (prepulse), presented approximately 20 to 500 ms before a startling sensory stimulus (pulse), inhibits the startle response to the startling stimulus (Graham, 1975; Norris and Blumenthal, 1996). Impaired PPI is observed in various psychiatric and neurological disorders, including schizophrenia, Huntington’s disease, Tourette’s syndrome, autism spectrum disorder, obsessive-compulsive disorder, post-traumatic stress disorder, bipolar disorder, seizure disorders, and nocturnal enuresis (Braff et al., 1978; Braff and Geyer, 1990; Swerdlow et al., 1995; Pouretemad et al., 1998; Ornitz et al., 1999; Perry et al., 2001; Baeyens et al., 2006; Gogos et al., 2009; Eggert et al., 2012; Holstein et al., 2013; Swerdlow, 2013; Schulz-Juergensen et al., 2014; Ahmari and Dougherty, 2015; Ahmari et al., 2016; Sánchez-Morla et al., 2016; Cavanna et al., 2017). Impaired PPI is often associated with symptoms common to these disorders, including psychosis, attention deficits, sensory overload, and other cognitive and motor symptoms.

While the reversal of PPI deficits in patients and animal models has been used as a tool for antipsychotic drug screening (Kumari et al., 1999; Swerdlow et al., 1999; Ribeiro et al., 2013), common antipsychotics show inconsistent effects (reviewed in Swerdlow and Light, 2016). This inconsistency is partly due to gaps in our knowledge of the neuronal pathways and mechanisms underlying the modulation of PPI. Therefore, identifying the cell types and synaptic mechanisms involved in PPI modulation is crucial and has potential clinical applications. Addressing this knowledge gap will improve our understanding of the pathophysiology of disorders associated with PPI deficits and facilitate the development and screening of therapeutics for these conditions.

While PPI can be induced by different sensory modalities, PPI of acoustic startle response is mostly used in clinical settings. The mammalian acoustic startle circuit is straightforward (**Fig. 1**, red pathway): in the brainstem, upon auditory stimulation, cochlear root neurons first activate contralateral giant neurons in the caudal pontine reticular nucleus (PnC). These startle-mediating giant PnC neurons then directly innervate cervical and spinal motor neurons, producing motor responses (Davis et al., 1982; Lee et al., 1996; Li et al., 2009). Inhibition of this startle pathway by non-startling prepulses at the level of the PnC leads to PPI (**Fig. 1**, blue pathway), via the cochlear nucleus and other midbrain and higher structures (Graham, 1975; Braff et al., 1978; Swerdlow et al., 1993; Perry and Braff, 1994; Castellanos et al., 1996; Grillon et al., 1996; Hoenig et al., 2005; Li et al., 2009). Previous animal and human studies have shown that the activation of midbrain structures, including the inferior and superior colliculi, as well as the pedunculopontine tegmental nucleus (PPTg) regulate PPI (Semba and Fibiger, 1992; Koch et al., 1993; Swerdlow and Geyer, 1993; Yeomans et al. 2006). Additionally, various cortical and subcortical areas within the cortico-striatal-pallido-pontine (CSPP) circuit such as the prefrontal cortex, thalamus, hippocampus, nucleus accumbens, and dorsal striatum, have been shown to affect PPI at different inter-stimulus intervals between prepulse and pulse (reviewed in Swerdlow et al., 2001; Fendt et al., 2001; and in Li et al., 2009).

**Figure 1.**
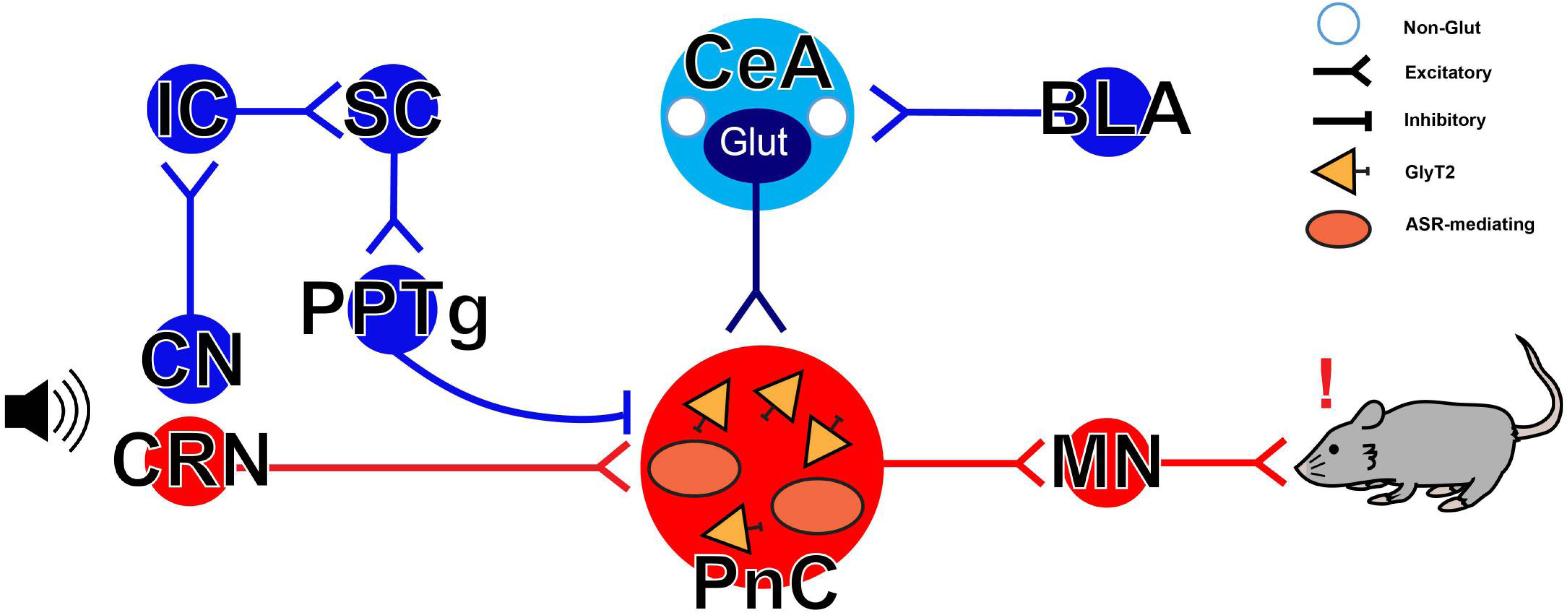
Simplified neuronal circuits underlying the acoustic startle response and PPI. The mammalian primary acoustic startle pathway (red pathway) includes primary auditory neurons that activate cochlear nuclei (CN) and cochlear root neurons (CRNs) which then transmit auditory information mostly to contralateral startle-mediating giant neurons of the brainstem caudal pontine reticular nucleus (PnC). These PnC giant neurons directly activate cervical and spinal motor neurons (MNs), leading to a motor response. During PPI, the startle pathway has been thought to be inhibited by acoustic prepulses via an ascending pathway (blue pathway) including the inferior (IC) and superior colliculi (SC), as well as the pedunculotegmental area (PPTg). We recently showed that the central nucleus of the amygdala (CeA) contributes to PPI by sending glutamatergic inputs to PnC neurons expressing the glycine transporter type 2 (GlyT2). BLA: basolateral amygdala; ASR: acoustic startle response. Modified from Cano et al., 2021.

There is also evidence showing the involvement of the amygdala in regulating PPI. Dys-functions in the amygdala are commonly associated with pathologies that exhibit sensorimotor gating deficits (Ludewig et al., 2002; Howland et al., 2007; Takeuchi et al., 2011; Rajbhandari et al., 2015; Wolf et al., 2015; Barth et al., 2021). The basolateral nucleus of the amygdala (BLA) receives inputs from multiple sensory systems (Sah et al., 2003; LeDoux, 2007), including auditory inputs, and projects to the central nucleus of the amygdala (CeA). The CeA, as the major output region of the amygdala, sends divergent projections to the basal forebrain, thalamus, hypothalamus, and brainstem (Sah et al., 2003; LeDoux, 2007; Moscarello and Penzo, 2022). Though mainly considered as a striatum-like structure composed of GABAergic cells resembling medium spiny neurons (McDonald and Augustine, 1993; Sun and Cassell, 1993; Pitkänen and Amaral, 1994; Saha et al., 2000), the CeA also sends glutamatergic projections to the PnC, as demonstrated in studies on guinea pigs (Zhang et al., 2012) and rodents (Rosen et al., 1991; Koch and Ebert, 1993; Lingenhöhl and Friauf, 1994). In fact, early stimulation studies showed that startle-mediating PnC giant neurons can be directly activated by excitatory CeA inputs with an average latency of 2.9 ms (Lingenhöhl and Friauf, 1994) in the context of fear-potentiated startle in rodents (Hitchcock and Davis, 1986; Rosen and Davis, 1988; Hitchcock and Davis, 1991; Hartley et al., 2019) and in human studies (Grillon et al., 1991).

Our recent mouse studies were the first to directly investigate the function of CeA glutamatergic neurons projecting to the PnC in the context of PPI (Cano et al., 2021; Huang et al., 2024), beyond their known role in fear-potentiated startle. Our anatomical and *in vitro* electrophysiological data, combined with optogenetic manipulations, demonstrated that CeA glutamatergic inputs can directly activate PnC neurons expressing the glycinergic transporter type 2 (GlyT2; Zeilhofer et al., 2005). We also functionally confirmed the involvement of CeA glutamatergic inputs and GlyT2^+^ PnC glycinergic neurons during PPI *in vivo* (Cano et al., 2021; Huang et al., 2024). These mechanistic findings are consistent with c-fos expression in the CeA observed during PPI in rats (Tapias-Espinosa et al., 2019). However, the specific functional networks involving CeA and PnC neuronal populations during acoustic startle and PPI remain to be fully elucidated. Furthermore, it is still unknown how these PPI-related functional networks are affected by disease states.

Here, we used the Cal-Light system to tag only neurons that are within the mouse CeA-PnC pathway and specifically involved in acoustic startle modulation with high spatiotemporal precision. Cal-Light is an innovative, calcium-dependent and light-sensitive tagging method that allows for labeling and optogenetic manipulation of neurons active during specific behavioral epochs (Lee et al., 2017; Hyun et al., 2022; **Suppl. Fig. 1**). Using Cal-Light, along with tract tracing, *in situ* hybridization, immunohistochemical, and cell counting analyses, we hypothesized that distinct neuronal populations in the CeA and PnC contribute to PPI and could serve as potential disease-relevant targets. We validated this hypothesis by optogenetically manipulating CeA-PnC glutamatergic synapses in *Prodh^−/−^* mice, a schizophrenia-relevant model exhibiting PPI deficits as well as anatomical and electrophysiological dysfunctions in the CeA.

## Results

### Identifying amygdala neurons active during PPI

Using *in situ* hybridization, immunohistochemistry, *in vitro* electrophysiological recordings, as well as *in vitro* and *in vivo* optogenetic manipulations in transgenic and wild-type (WT) mice, we previously demonstrated that CeA neurons send glutamatergic inputs to the PnC and contribute to PPI (Cano et al., 2021; Huang et al., 2024). While our results established a direct circuit-behavior relationship, selectively labeling and manipulating CeA neurons active during PPI is critical to confirm their causal role.

Until now, the specific identification of PPI-related neurons has been challenging due to the difficulty of labeling neurons involved in fast, transient, and short-lived behaviors like PPI. To overcome this limitation and selectively label PPI-specific neurons on a very short time scale, we unilaterally injected Cal-Light viral components into the CeA of WT mice (N = 5; **Suppl. Fig. 1 and Fig. 2**). We then implanted a cannula in the CeA to position an optic fiber for light delivery dorsal to the Cal-Light-targeted neurons. Neurons expressing Cal-Light should exhibit an initial red fluorescent signal through tdTomato expression within approximately 5-10 days (Lee et al., 2017; Hyun et al., 2022). When these Cal-Light^+^ neurons are activated, the resulting rise in intra-cellular calcium combined with exposure to blue light (470 nm) should lead to the expression of eGFP together with the yellow light-sensitive inhibitory optogenetic tool halorhodopsin (NpHR3.0) within 2-5 days.

**Figure 2.**
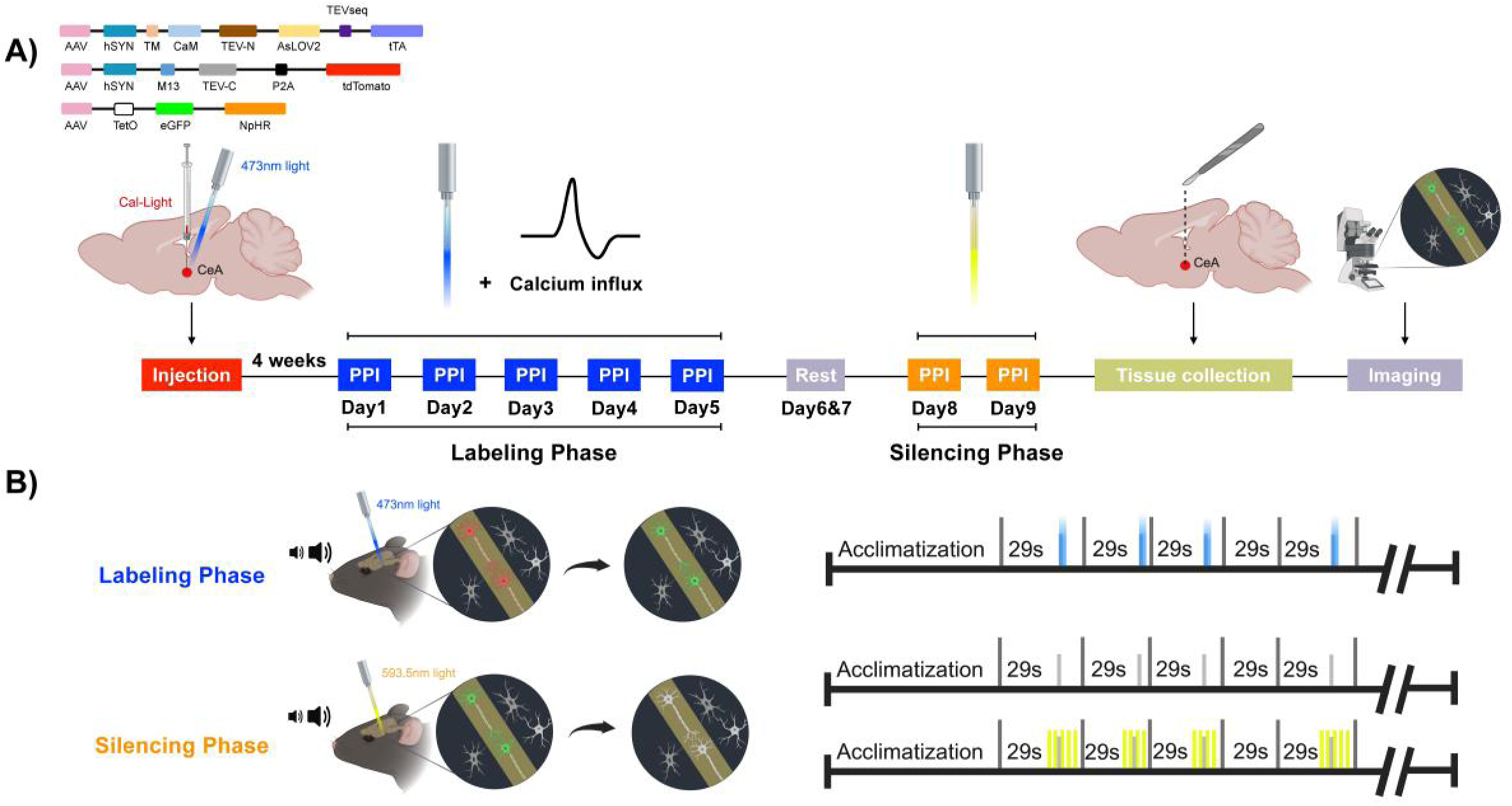
Photo labeling and silencing Cal-Light^+^ CeA neurons active during PPI. **A)** Behavioral protocol used to photo-label CeA neurons active during PPI. Cal-Light viruses including eGFP and NpHR reporters were unilaterally injected into the CeA of WT mice to specifically label and then inhibit CeA neurons active during PPI. Four weeks after the viral injection, all mice underwent a 5-day labeling phase followed by a 2-day resting phase and then a 2-day silencing phase. After the silencing phase, the brain was harvested and thin CeA sections were obtained to visualize active Cal-Light-targeted CeA neurons expressing eGFP. **B)** During the 5-day labeling phase, blue light was delivered concurrent with the prepulses of PPI trials. This made CeA neurons active during PPI express eGFP and NpHR. This phase was followed by a resting period. Then, during the silencing phase, the blue light was replaced by a yellow light delivered to inhibit eGFP and NpHR expressing CeA neurons during the prepulses of PPI trials.

First, we wanted to visualize CeA neurons active during PPI using eGFP expression. We hypothesized that these neurons would be activated by a prepulse and remain active during the interval between the prepulse and the pulse (i.e., inter-stimulus interval or “ISI”) of PPI trials. Therefore, to label or photo-tag neurons active during PPI, blue light was delivered to CeA neurons only during the prepulses and inter-stimulus intervals (**Fig. 2**), for a total of 9 s across all trials in Cal-Light-injected mice. To maximize eGFP labeling of active neurons, the blue light photo-tagging phase was performed over five consecutive days, resulting in robust eGFP expression (**Fig. 2 and Fig. 3**). When we quantified the number of eGFP^+^ CeA neurons, our results showed that 68.45% ± 1.78% of Cal-Light-targeted neurons were active during PPI at all intervals tested and expressed eGFP. Then, to chemically identify these PPI-related Cal-Light/eGFP^+^ CeA neurons, we performed *in situ* hybridization using the RNAscope assay. Our analysis revealed that 92.27% ± 9.25% eGFP^+^ CeA neurons expressed the vesicular glutamate transporter type 2 (i.e., VGLUT2^+^; **Fig. 4**, N = 4).

**Figure 3.**
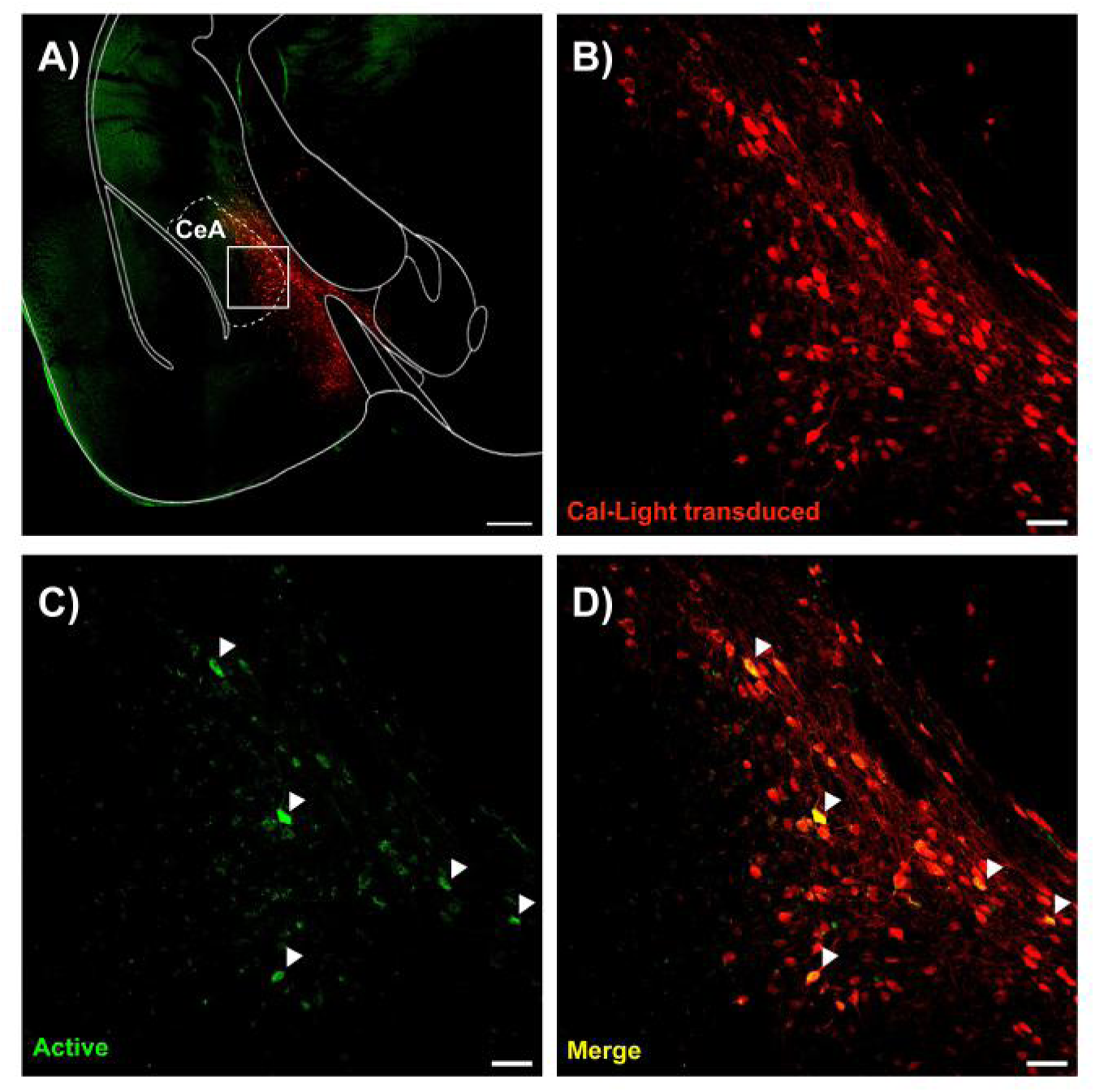
Cal-Light^+^ CeA neurons active during PPI. **A)** Low magnification of CeA neurons targeted with Cal-Light and expressing the transduction marker TdTomato (red) and the Cal-Light reporter eGFP. The CeA is delineated by dotted lines. The white rectangle shows the area imaged in panels B-D. **B)** CeA neurons injected with Cal-Light expressed tdTomato. **C)** In the presence of blue light, CeA neurons active during PPI expressed eGFP. **D)** Merged image showing that only a subset of Cal-Light targeted CeA neurons were active during PPI. Arrowheads indicate active CeA neurons expressing both TdTomato and eGFP. Scale bars: A, 250μm; B-D, 50μm. Representative of N = 5 WT mice.

**Figure 4.**
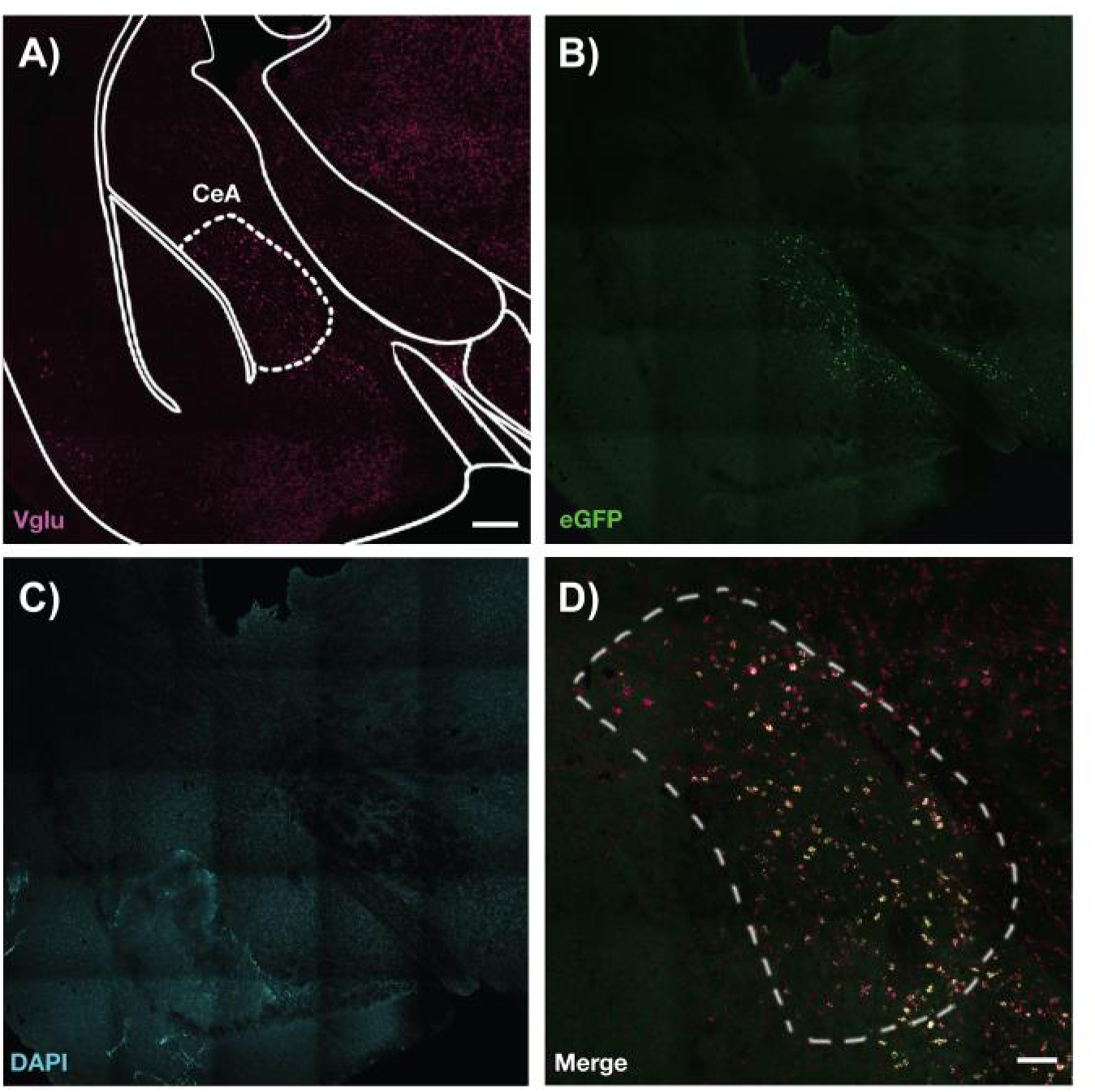
Cal-Light^+^ CeA neurons active during PPI are VGluT2 positive. **A-C)** Representative images from Cal-Light injected mice following the labeling and silencing phases. The CeA coronal section was hybridized with VGluT2 (magenta) and eYFP (green) probes and stained with DAPI (cyan) to label cell bodies. The CeA is delineated by dotted lines. **D)** Higher magnification and merged image of the CeA area in panels A-C, where white cells represent Cal-Light targeted and VGluT2^+^ CeA neurons active during PPI. Representative of N = 4 WT mice. Scale bars: A-C, 250μm; D, 50μm.

Next, we wanted to test whether these Cal-Light/eGFP^+^ CeA neurons contribute to PPI. In Cal-Light-injected mice, following the blue-light photo-tagging phase described above, CeA neurons active during PPI would express both eGFP and NpHR3.0. This allowed us to use yellow light (589 nm) to photo-inhibit these eGFP/NpHR3.0^+^ CeA neurons during startle and PPI (**Fig. 2 and Fig. 5**). We hypothesized that altering neuronal activity of these neurons would perturb PPI without affecting baseline startle. We first confirmed that in the absence of yellow light, Cal-Light-injected mice exhibited measurable acoustic startle reflexes, characterized by a whole-body flexor muscle contraction (Davis et al., 1982; Lee et al., 1996; Swerdlow et al., 1999), when reacting to acoustic pulses above 90 dB. Then, we tested whether silencing eGFP/NpHR3.0^+^ CeA glutamatergic neurons active during PPI would affect baseline startle. To do so, we photo-inhibited these neurons with yellow light during acoustic pulse-alone stimulations at increasing sound levels, and observed startle amplitudes comparable to those measured without yellow light (light effect: F_(1,8)_ = 0.17, *p* = 0.691; **Fig. 5A, B**). These results suggest that CeA glutamatergic neurons active during PPI do not contribute to the baseline startle response. Then, we tested whether eGFP/NpHR3.0^+^ CeA glutamatergic neurons contribute to PPI. Our results show that photo-inhibiting these CeA neurons during the prepulse significantly reduced PPI by 40-50% at ISIs between 10-50 ms (light effect, F_(1,24)_ = 39.35, *p* < 0.001; Two-way RM ANOVA; **Fig. 5C, D**). This is aligned with our previous findings through the optogenetic inhibition of CaMKIIα/NpHR3.0^+^ CeA glutamatergic neurons during the prepulse in WT mice, that also led to reduced PPI at short ISIs of 30 and 50 ms (Cano et al., 2021). Taken together, our data confirms that these eGFP/NpHR3.0^+^ CeA glutamatergic neurons contribute to PPI at short interstimulus intervals between the prepulse and pulse.

**Figure 5.**
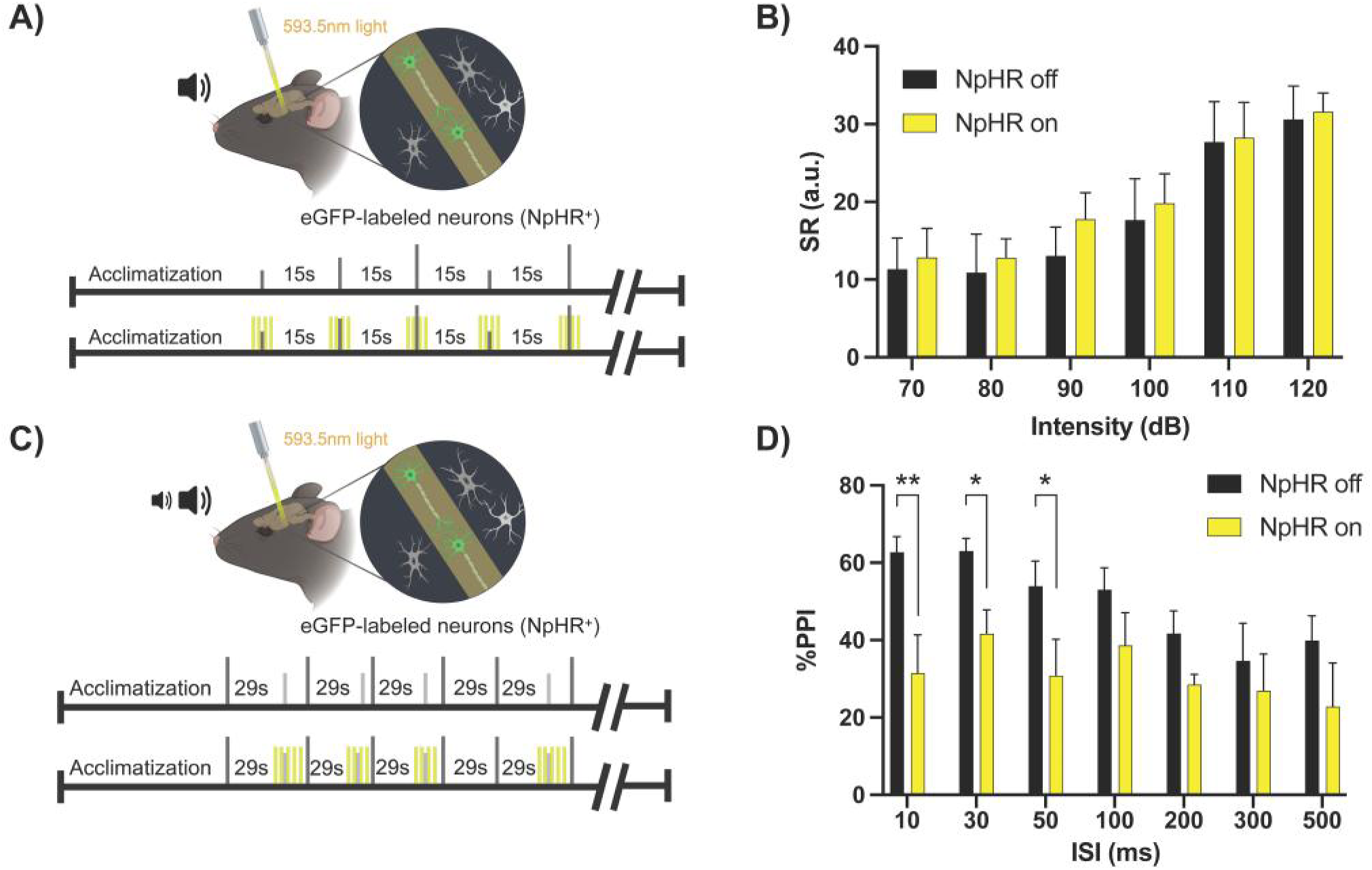
Photo-inhibition of Cal-Light^+^ CeA neurons active during PPI. **A)** Schematic of the acoustic startle response protocols performed in WT mice injected with Cal-Light viral components in the CeA. In these mice, Cal-Light-targeted CeA neurons active during PPI expressed the yellow light-sensitive inhibitory optogenetic tool, NpHR. **B)** Graph showing the mean startle response (SR) amplitude as a function of sound at increasing intensities, in the absence (black bars; “NpHR off”) and in the presence (yellow bars; “NpHR on”) of yellow light delivered concurrent with acoustic pulses. Photo-inhibition of Cal-light targeted CeA neurons active during PPI did not affect the amplitude of baseline startle elicited by 70-120 dB pulse alone acoustic stimulations [light effect: F_(1,8)_ = 0.17, *p* = 0.691]. **C)** Schematic of the PPI protocols performed in WT mice injected with Cal-Light viral components in the CeA. **D)** Graph showing mean PPI values as a function of interstimulus intervals (ISI) between acoustic prepulse and pulse, in the absence (black bars; “NpHR off”) and in the presence (yellow bars; “NpHR on”) of yellow light delivered concurrent with acoustic prepulses. Photo-inhibition of Cal-Light-targeted CeA neurons active during PPI, significantly decreased PPI at shorter ISIs in WT mice [light effect, F_(1,24)_ = 39.35, *p* < 0.001]. Two-way RM ANOVA, N = 5 WT mice. Data are represented as mean ± SEM. \**p* < 0.05, \*\**p* < 0.01.

Overall, these results demonstrate that the Cal-Light system can precisely tag neurons activated on a millisecond timescale with eGFP and NpHR in specific brain regions. This allowed us to selectively label and photo-inhibit PPI-related neurons in the CeA, offering critical insight into the timing and extent of their contribution to PPI.

### Identifying PnC neurons active during acoustic startle and PPI

The PnC lies at the intersection of the acoustic startle and PPI pathways (**Fig. 1**), playing a key role in mediating both behavioral responses. It is known to contain a heterogeneous cellular population which includes small inhibitory neurons interspersed among startle-mediating PnC giant neurons (Davis et al., 1982; Lingenhöhl and Friauf, 1992, 1994; Lee et al., 1996). However, the specific identity of PnC neurons active during startle versus PPI remains unclear. Therefore, we aimed to identify the distribution and neurochemistry of PnC neurons active during acoustic startle and PPI. Similar to the approach we used to identify active CeA neurons, we next injected Cal-Light viral components into the PnC of WT mice (**Fig. 6**; N = 11) and unilaterally implanted an optic fiber for blue light delivery.

**Figure 6.**
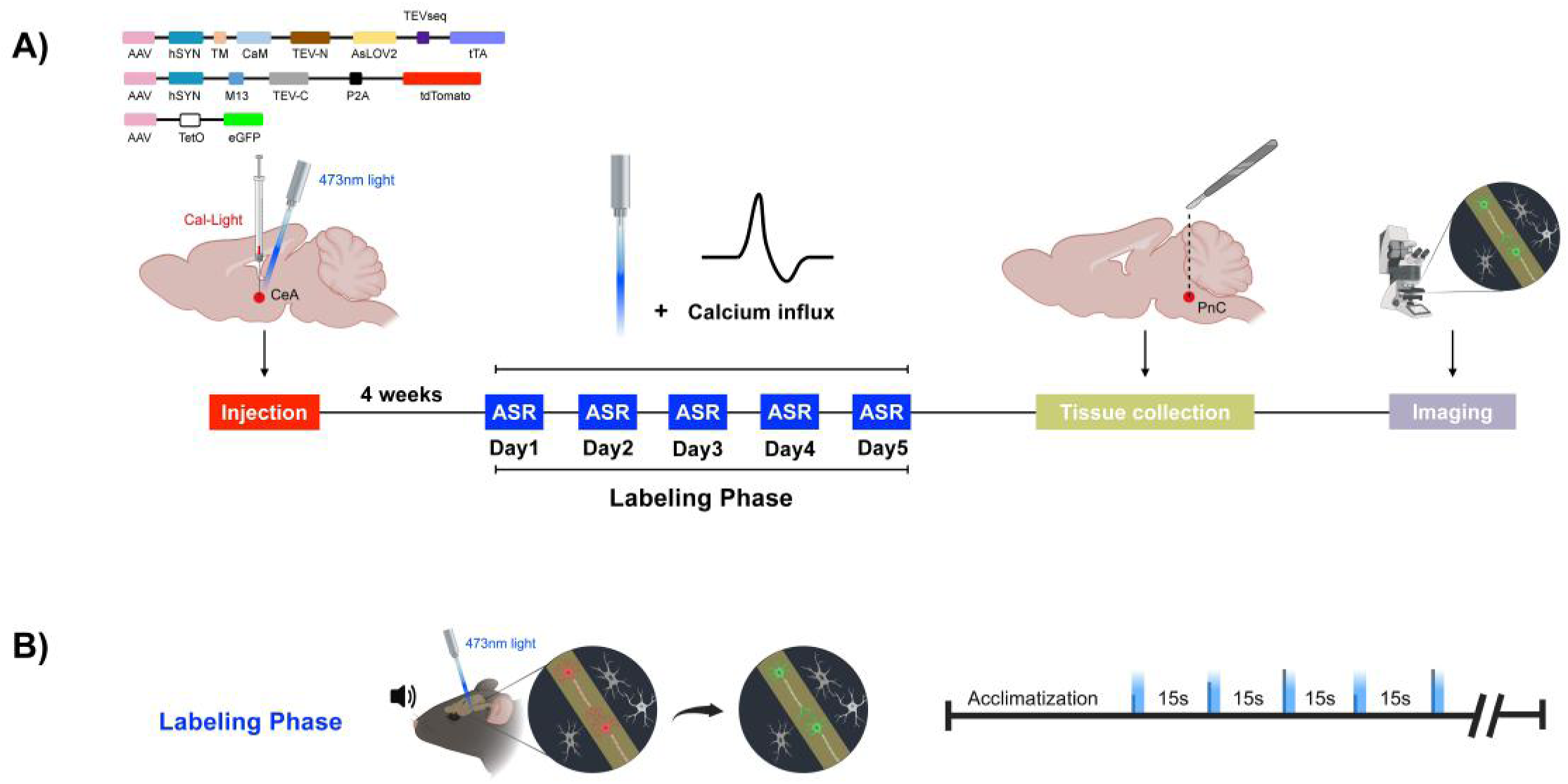
Using Cal-Light to label PnC neurons active during the acoustic startle response. **A)** Behavioral protocols used to label PnC neurons active during the acoustic startle response (ASR). Cal-Light viruses were injected into the PnC of WT mice unilaterally. Acoustic stimuli were delivered four weeks after the injection to elicit baseline startle responses. Following the labeling phase, the brain was harvested and thin PnC sections were obtained to visualize Cal-Light targeted neurons active during baseline startle with eGFP. **B)** All mice underwent a 5-day labeling phase during which, blue light was delivered concurrent with the acoustic pulses of ASR trials. This made Cal-Light targeted PnC neurons active during startle express eGFP.

To label PnC neurons active during startle responses, blue light was delivered to Cal-Light^+^ PnC neurons during startling pulses (120 dB, 40 ms; **Fig. 6**) of startle trials. All PnC neurons transduced with Cal-Light were imaged, focusing specifically on the subset of active neurons expressing eGFP (**Suppl. Fig. 2**). The density and cell types of PnC neurons active during startle were characterized by staining PnC sections across six different anatomical levels (mapped on levels 75-78 of the Paxinos and Franklin Mouse Brain Atlas, 2004) with an antibody for parvalbumin (PV) to quantify PV^+^ inhibitory neurons (**Fig. 7**). Our cell counting analysis revealed that, among the Cal-Light^+^ PnC neurons transduced with tdTomato (**Fig. 7A**), 50.68% ± 3.09% were active and expressed eGFP during the startle assay (**Fig. 7B**), out of which 14.93% ± 2.71% were also PV^+^ (N = 4, **Fig. 7C-E**). This indicates that apart from excitatory startle-mediating giant neurons, a small subset of PV^+^ inhibitory neurons also contributes to startle responses in the PnC.

**Figure 7.**
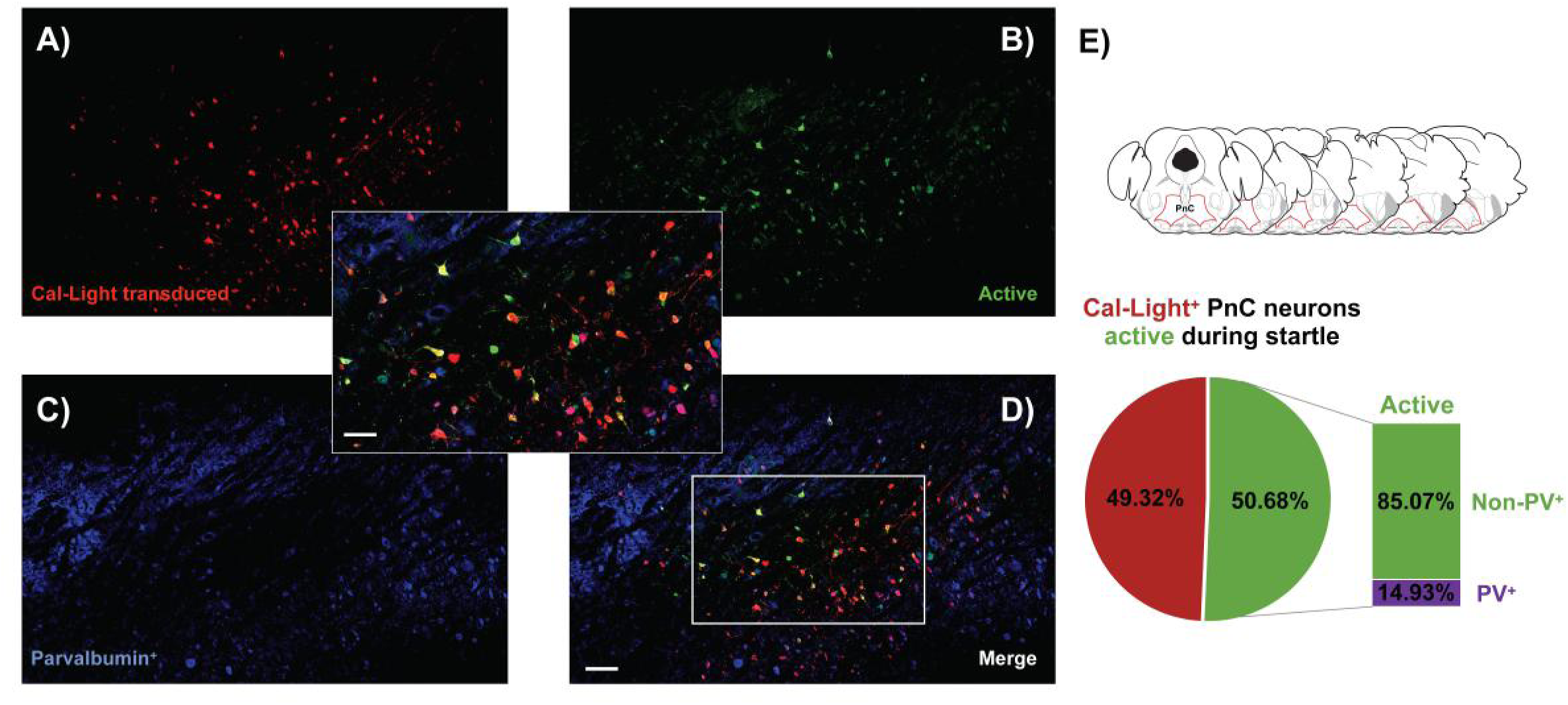
A subset of Cal-Light^+^ PnC neurons active during baseline startle are inhibitory. **A)** PnC neurons targeted with Cal-Light and expressing the transduction marker TdTomato (red). **B)** A subset of Cal-Light**^+^** PnC neurons that were active during baseline startle, and expressing the reporter eGFP (green) in the presence of blue light. **C)** PnC neurons co-labeled with the parvalbumin (PV) antibody. **D)** Merged image showing that a few Cal-Light-targeted PnC neurons active during baseline startle were PV**^+^**. ***Inset***, Boxed area in panel D shown at higher magnification. **E)** Ratio of inhibitory PnC neurons active during baseline startle. Nearly 51% of Cal-Light^+^ neurons were active during startle, out of which 15% were PV^+^. Cell counting was performed across six different anatomical levels of the PnC, mapped onto the Paxinos and Franklin Mouse Brain Atlas. Scale bars: A-D, 100μm; Inset, 50μm. Representative of N = 4 WT mice.

Next, we labeled PnC neurons active during PPI by delivering blue light during the prepulses (75 dB, 20 ms; **Fig. 8**) and during the entire inter-stimulus intervals of PPI trials. Interestingly, our results show that among all the neurons transduced with Cal-Light and expressing tdTomato (**Fig. 9A**), 68.59% ± 13.74% were active during PPI and expressed eGFP (**Fig. 9B**), out of which 24.58% ± 2.71% were PV^+^ (N = 4, **Fig. 9 C-E**). These data suggest that a greater proportion of PnC inhibitory neurons are active during PPI than during startle alone. This is consistent with our previous findings that photo-manipulation of inhibitory glycinergic neurons co-expressing PV only affects PPI without altering startle amplitudes (Cano et al., 2021; Huang et al., 2024). Taken together, Cal-Light enabled us to identify a specific subset of PV^+^ PnC neurons active during startle and PPI, clearly highlighting the involvement of PnC inhibitory networks during PPI.

**Figure 8.**
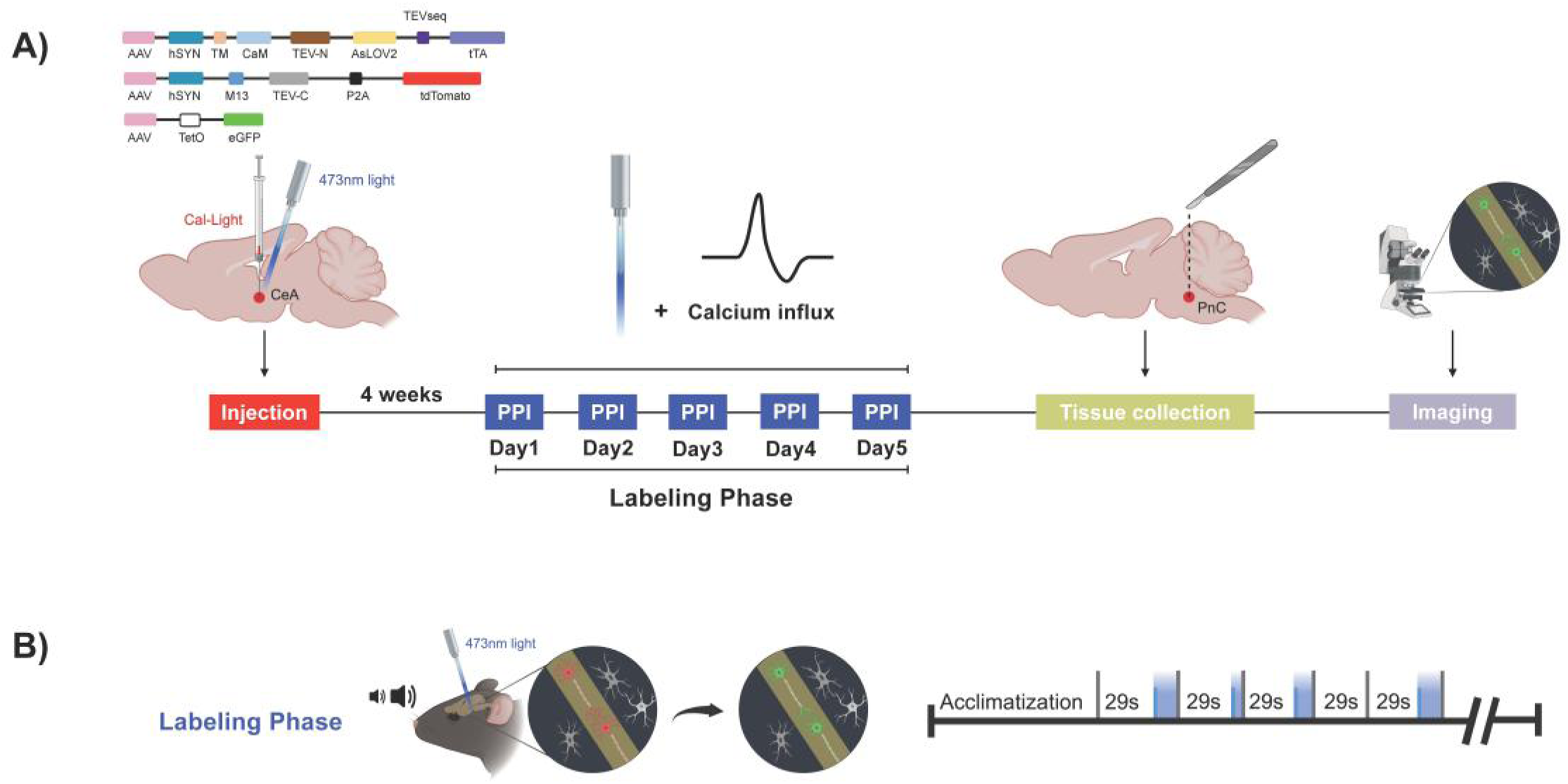
Using Cal-Light to label PnC neurons active during PPI. **A)** Behavioral protocols used to label PnC neurons active during PPI. Cal-Light viruses were injected into the PnC of WT mice unilaterally. Acoustic stimuli were delivered four weeks after the injection to elicit PPI. Following the labeling phase, the brain was harvested and thin PnC sections were obtained to visualize Cal-Light targeted neurons active during PPI with eGFP. **B)** All mice underwent a 5-day labeling phase during which, blue light was delivered concurrent with the acoustic prepulses and interstimulus intervals of PPI trials. This made Cal-Light-targeted PnC neurons active during PPI express eGFP.

**Figure 9.**
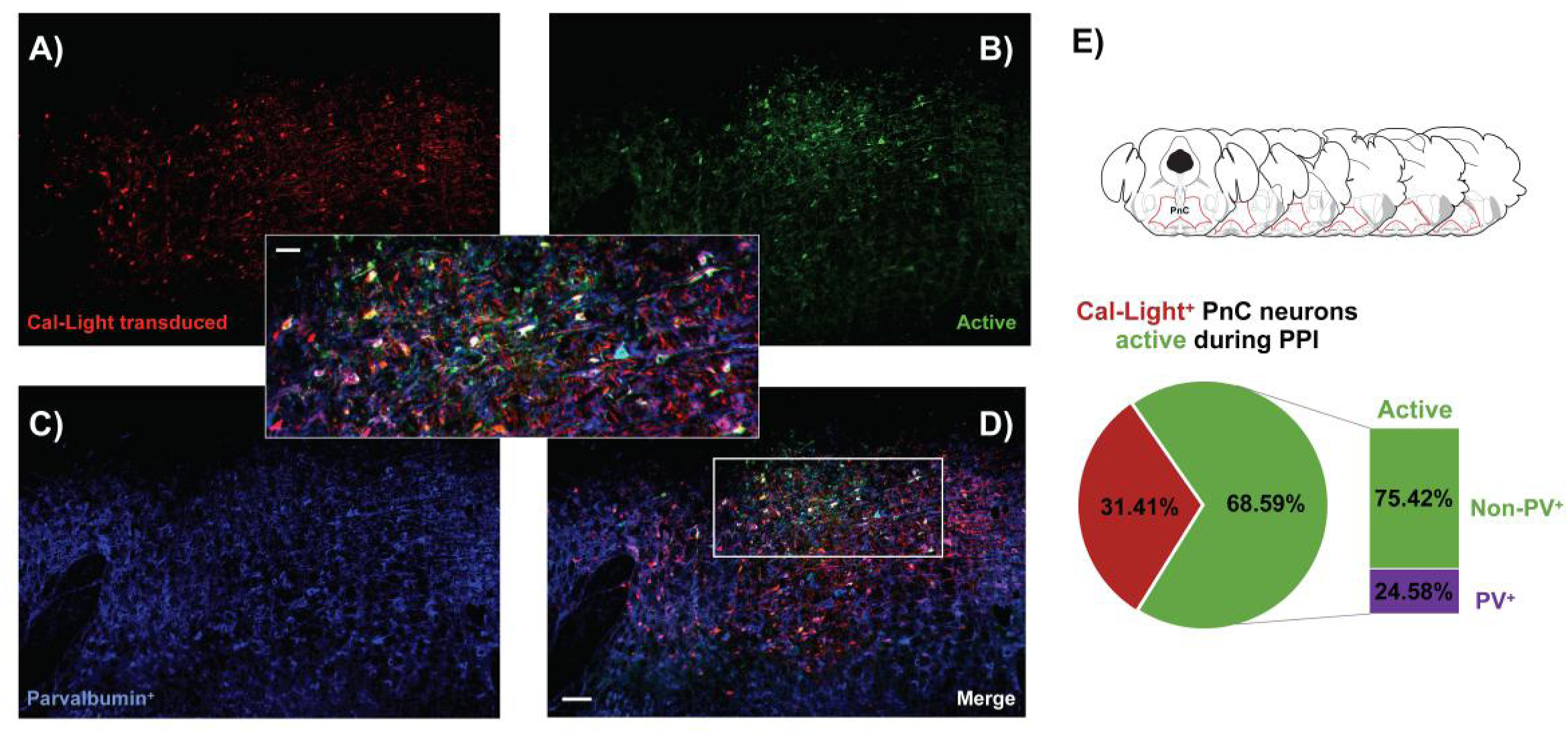
The ratio of active Cal-Light^+^ PnC inhibitory neurons increases during PPI. **A)** PnC neurons targeted with Cal-Light and expressing the transduction marker TdTomato (red). **B)** A subset of Cal-Light**^+^** PnC neurons that were active during PPI, and expressing the reporter eGFP in the presence of blue light. **C)** PnC neurons co-labeled with the parvalbumin antibody. **D)** Merged image showing that several Cal-Light-targeted PnC neurons active during PPI were PV^+^. ***Inset***, Boxed area in panel D shown at higher magnification. **E)** The ratio of active PnC inhibitory neurons increases during PPI. Nearly 69% of Cal-Light^+^ neurons were active during PPI, out of which 25% were PV^+^. Cell counting was performed across six different anatomical levels of the PnC, mapped onto the Paxinos and Franklin Mouse Brain Atlas. Scale bars: A-D, 100μm; Inset, 50μm. Representative of N = 4 WT mice.

Lastly, we performed control experiments to confirm that the Cal-Light system only labels active PnC neurons exposed to blue light. We injected Cal-Light components into the PnC of WT mice (N = 12) and unilaterally implanted an optic fiber for blue light delivery. One group of injected mice were freely moving in a sound-attenuated chamber with constant background noise, either in the absence of (**Suppl. Fig. 3**, “no light, no activity”; N = 4) or in the presence of blue light exposure (the same light duration as used for PPI trials; **Suppl. Fig. 3**, “light only”; N = 4). In another group, injected mice underwent the same PPI trials, but without blue light exposure (**Suppl. Fig. 3**, “activity only”; N = 4). Next, we examined the brains of these injected mice. In the PnC of these mice, we detected robust tdTomato expression but minimal eGFP expression, confirming that Cal-Light is activity and blue light-dependent. Overall, these data show that only mice both exposed to blue light and reactive to acoustic startle stimulations or PPI had neurons expressing robust eGFP in the PnC.

### Photo-activating CeA-PnC glutamatergic synapses restores PPI in mice with Proline dehydrogenase (*PRODH*) deficiency

*PRODH*, the gene encoding the proline dehydrogenase enzyme, resides within the schizo-phrenia-linked 22q11.2 locus and functions to degrade L-proline. Deletion or dysfunction of *PRODH* leads to elevated CNS L-proline levels, which are associated with an increased risk of psychotic disorders (Raux et al., 2007; Willis et al., 2008; Clelland et al., 2011), including schizophrenia. *Prodh^−/−^* mice (transgenic mice with mutations in both copies of the *Prodh* gene) show increased proline levels in PPI-related brain regions, including the amygdala (Gogos et al., 1999). Additionally, *Prodh^−/−^* mice display behavioral deficits reminiscent of those seen in schizophrenia patients, including impaired fear conditioning and reduced PPI, both of which depend on proper amygdala function (Gogos et al., 1999). Therefore, we used *Prodh^−/−^*mice to validate that the CeA-PnC connection is a site of disease-related PPI dysfunction.

Before investigating the potential impact of *Prodh* deficiency at the brainstem CeA-PnC connection, we first assessed the expression of *Prodh* in PnC neurons. In the medial prefrontal cortex (mPFC) of WT mice, *Prodh* is normally widely expressed in neurons, including PV^+^ inhibitory neurons (Crabtree et al., 2016), which comprise both GABAergic and glycinergic neurons. Since *Prodh^−/−^*mice exhibit altered inhibitory neurotransmission in the mPFC, we hypothesized that *Prodh* is also expressed in PnC inhibitory neurons and that its dysfunction would impact PnC inhibitory signaling. To test this hypothesis, we injected a Cre-dependent eYFP neuronal tracer into the PnC of *GlyT2^Cre^* mice, where Cre is expressed under the control of the glycine transporter type 2 (GlyT2) promoter (N = 5). Our immunohistochemical analyses, using PRODH and PV antibodies, show that PRODH is co-expressed with PV and eYFP in GlyT2^+^ PnC neurons (**Suppl. Fig. 4**)

Next, we assessed how *Prodh* deficiency impacts PnC inhibitory signaling during startle and PPI. We first conducted *in vivo* behavioral testing on *Prodh^−/−^* mice (N = 9; 2-3 months old) and their WT littermates (N = 6; 2-3 months old). Consistent with previous studies (Gogos et al., 1999), we found that *Prodh^−/−^* mice exhibited reduced PPI compared to WT controls (Main effect of genotype: F_(1,15)_ = 10.01, *p* = 0.006; Mixed ANOVA; **Fig. 10 A-C**). To determine whether the reduced PPI in *Prodh^−/−^* mice was attributable to changes in baseline startle reactivity, we compared the amplitude of startle responses between genotypes. As expected, there was no change in baseline startle reactivity between genotypes (Main effect of genotype: F_(1,13)_ = 1.564, *p* = 0.233; **Suppl. Fig. 5**), suggesting that the reduced PPI levels observed in *Prodh^−/−^* mice are not correlated with changes in baseline startle reactivity. In other words, *Prodh* deficiency only affects PPI without altering startle responses.

**Figure 10.**
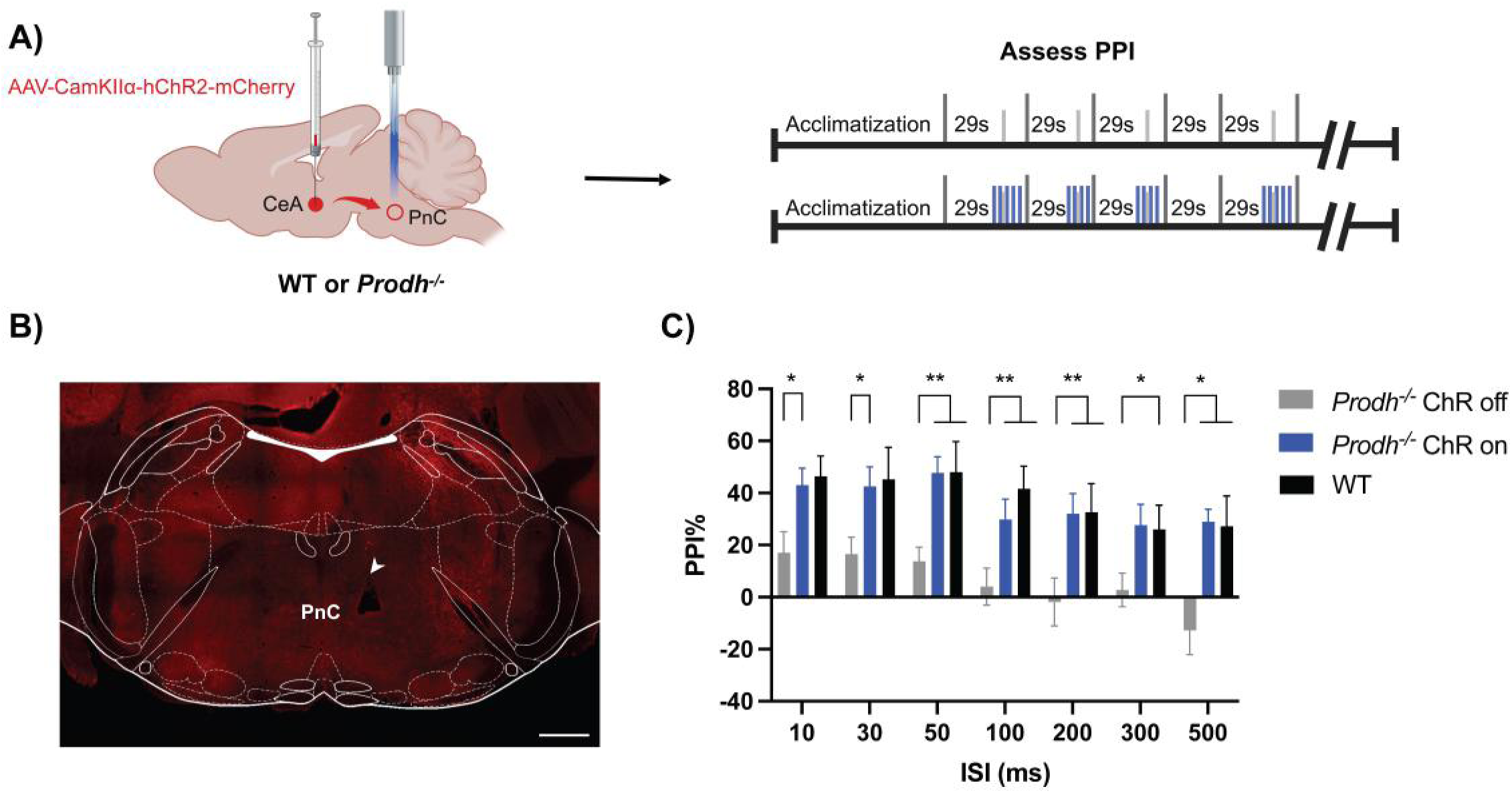
Photo-activation of CeA-PnC glutamatergic synapses rescues PPI deficits in *Prodh^−/−^* mice. **A) *Left***, Schematic illustration of the mouse brain sagittal section showing the injection site of the anterograde AAV-DJ-CamKIIα-hChR2-mCherry virus in the CeA and the optic fiber implanted unilaterally in the PnC of *Prodh^−/−^* or WT mice. ***Right***, Schematic of PPI protocols performed in mice injected with ChR2. **B)** Representative image of a coronal PnC section delineated by the 7^th^ cranial nerves showing the cannula tract (white arrowhead). **C)** Graph showing mean PPI values as a function of interstimulus intervals (ISIs) between acoustic prepulse and pulse, both in the absence (grey bars; “ChR off”) and in the presence (blue bars; “ChR on”) of blue light photo-activation of CeA-PnC glutamatergic synapses in *Prodh^−/−^* mice. Baseline PPI values in WT mice without light manipulation are shown in black bars. *Prodh^−/−^* mice had reduced PPI compared to WT mice [group effect, F_(1,15)_ = 10.01, *p* = 0.006, Mixed ANOVA]. However, activation of CeA-PnC glutamatergic synapses during acoustic prepulses increased PPI in *Prodh^−/−^* mice to the level that can be comparable to WT mice. [*Prodh^−/−^* : ChR on vs ChR off, light effect, F_(1,18)_ = 14.20, *p* = 0.001, Two-way RM ANOVA; *Prodh^−/−^* ChR on vs WT, group effect, F_(1,13)_ = 0.046, *p* = 0.8338, Mixed ANOVA]. N = 9 *Prodh^−/−^* mice and N = 6 WT mice. Scale bar in B: 500 μm. Data in C are represented as mean ± SEM. \**p* < 0.05, \*\**p* < 0.01.

We then aimed to rescue the reduction in PPI of *Prodh^−/−^*mice by targeting the brainstem CeA-PnC connection. We previously demonstrated that photo-stimulating CeA glutamatergic inputs can activate GlyT2^+^ neurons in the PnC (Cano et al., 2021) and enhance PPI (Huang et al., 2024). Therefore, we anticipated that activating PnC-projecting CeA glutamatergic inputs would likely increase PnC inhibitory neurotransmission in *Prodh^−/−^*mice, thereby restoring their PPI. To do so, we transduced CeA glutamatergic cells with Channelrhodopsin-2 (ChR2, an excitatory optogenetic tool sensitive to blue light) by injecting the viral vector AAV-CaMKIIa-hChR2(H134R)-mCherry into the CeA of *Prodh^−/−^* mice (N = 9; **Fig. 10 A**). Surprisingly, our results show that in *Prodh^−/−^* mice, photo-activating CeA-PnC synapses with blue light trains (5 Hz, 3 ms pulses) during prepulses significantly increased PPI levels at all ISIs tested (**Fig. 10 C**; *Prodh^−/−^* “ChR on” vs “ChR off”, light effect, F_(1,18)_ = 14.20, *p* = 0.001, Two-way RM ANOVA). Notably, the photo-stimulation in *Prodh*⁻/⁻ mice increased PPI to levels indistinguishable from those of their WT littermates (**Fig. 10 C**; *Prodh^−/−^* “ChR on” vs WT, group effect, F_(1,13)_ = 0.046, *p* = 0.8338, Mixed ANOVA). Overall, our results indicate that activating CeA-PnC glutamatergic synapses fully rescues PPI levels in *Prodh^−/−^* mice.

To further determine whether activating CeA-PnC glutamatergic synapses alone is sufficient to induce PPI in *Prodh^−/−^* mice, we tested the effect of photo-activating these synapses prior to an acoustic startle stimulation, in the absence of an acoustic prepulse. CeA-PnC glutamatergic synapses were photo-activated with blue light pulses (3 pulses at 50 Hz, 15 ms ON, 5 ms OFF; **Suppl. Fig. 6**) at inter-stimulus intervals (ISIs) typically used for acoustic prepulses. Our results show that photo-activating CeA-PnC glutamatergic synapses *in lieu of* an acoustic prepulse successfully induced a WT-like acoustic PPI effect in *Prodh^−/−^* mice at all ISIs tested (Main effect of prepulses: F_(1,13)_ = 2.519, *p* = 0.137; **Suppl. Fig. 6**). These findings suggest that in *Prodh^−/−^* mice, activation of CeA-PnC glutamatergic synapses alone can suppress a subsequent startle response and induce WT-like acoustic PPI levels.

### *Prodh* deficiency affects the structure and electrical properties of PnC-projecting CeA neurons

To provide mechanistic insights into how schizophrenia-relevant *Prodh* deficiency affects PPI, we conducted neuroanatomical and *in vitro* electrophysiological experiments. First, to investigate whether *Prodh* deficiency alters the morphology of PnC-projecting CeA glutamatergic neurons, we labeled these neurons by injecting AAV-CamKIIα-mCherry into the CeA of *Prodh^−/−^* (N = 9) and WT mice (N = 6). Interestingly, compared to WT littermates, CeA glutamatergic fibers of *Prodh^−/−^* mice were significantly denser in the ventrolateral region of the PnC (**Suppl. Fig. 7**).

Next, to determine whether these morphological changes in PnC-projecting CeA glutamatergic neurons are associated with altered electrical properties, we characterized both synaptic and intrinsic properties of these CeA neurons using whole-cell voltage-clamp recordings in *Prodh^−/−^* (N = 8) and WT mice (N = 7) injected with the retrograde AAVrg-CamKIIα-mCherry tracer into the PnC. Regarding postsynaptic properties, the kinetics of spontaneous excitatory postsynaptic currents (sEPSC) did not significantly differ between genotypes (**Suppl. Fig. 8 and Suppl. Table 1**): sEPSC rise time (ms) was WT = 0.80 ± 0.02 vs. *Prodh^−/−^* = 0.72 ± 0.05, while sEPSC decay time (ms) was WT = 4.1 ± 0.89 vs. *Prodh^−/−^* = 3.12 ± 0.54. Similarly, no significant differences were observed in EPSC frequency (#sEPSCs/min: WT = 487.9 ± 170.4 vs. *Prodh^−/−^* = 689.9 ± 290.9), amplitude (pA: WT = 9.7 ± 1.0 vs. *Prodh^−/−^* = 9.5 ± 1.2), or charge (FC: WT = 20.7 ± 3.4 vs. *Prodh^−/−^* = 25.4 ± 5.1).

Some of the intrinsic properties of these CeA neurons also appeared unaffected by *Prodh* deficiency (**Suppl. Table 1**). Capacitance values (pF) were WT = 19.0 ± 2.0 vs. *Prodh^−/−^* = 18.8 ± 2.0, series resistance (R_Series_, MΩ) was WT = 16.9 ± 2.5 vs. *Prodh^−/−^* = 18.5 ± 1.7, membrane resistance (*R_membrane_*, GΩ) was WT = 2.87 ± 0.80 vs. *Prodh^−/−^*= 4.56 ± 1.87, and input resistance (*R_Input_*, GΩ) was WT = 539.7 ± 34.0 vs. *Prodh^−/−^* = 562.22 ± 77.5. However, while the frequency of spontaneous action potentials (APs/min: WT = 27.8 ± 15.8 vs. *Prodh^−/−^* = 23.9 ± 12.6) and the action potential half-width at half-height (ms: WT = 1.49 ± 0.09 vs. *Prodh^−/−^* = 1.58 ± 0.08) were similar between genotypes, *Prodh^−/−^* mice exhibited a significantly lower rheobase compared to WT littermates (rheobase, pA: WT = 17.8 ± 6.62 vs. *Prodh^−/−^* = −6.1 ± 9.3). Interestingly, in *Prodh^−/−^* mice, this lower rheobase was associated with higher excitability quantified as higher AP firing rates upon current injection in PnC-projecting CeA glutamatergic neurons (**Fig. 11**).

**Figure 11.**
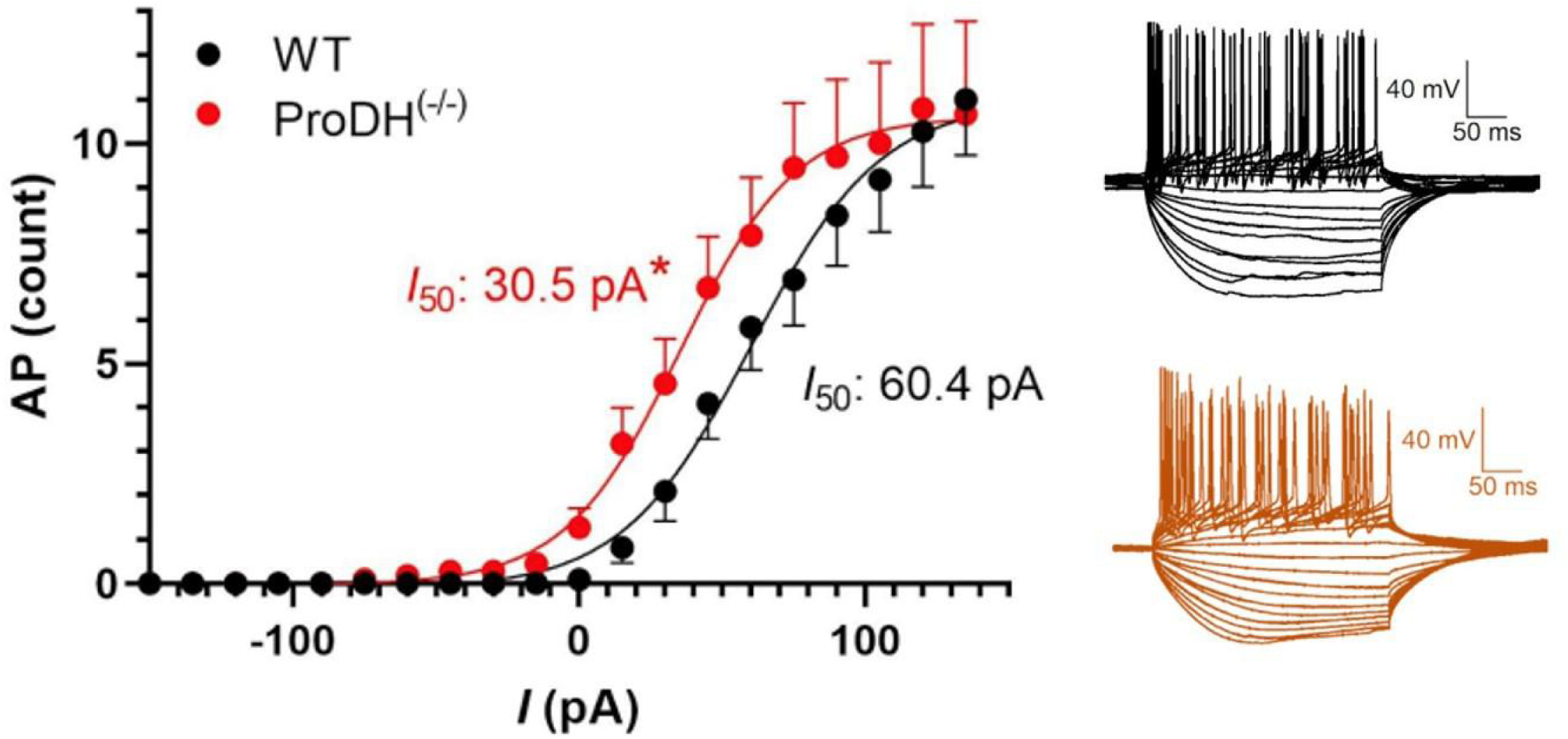
*Prodh* deficiency alters the firing properties of CamKIIα-ChR2 eYFP^+^ CeA glutamatergic neurons projecting to the PnC. Plot of the action potential (AP) firing rate as a function of depolarizing current injections. PnC-projecting CeA glutamatergic neurons of *Prodh^−/−^* mice (N = 8) display greater firing rates compared to WT mice (N = 7; *p* < 0.05; unpaired t-Test). ***Inset***, Representative voltage traces. Representative of n = 12 cells/genotype.

Overall, these neuroanatomical and electrophysiological data suggest that while CeA glutamatergic neurons display similar excitatory synaptic inputs, soma size, or number of open channels at rest between WT and *Prodh^−/−^*mice, these neurons are significantly more excitable in *Prodh^−/−^* mice. They also exhibit more axonal branching in the PnC. Such structural and functional changes at CeA-PnC synapses, alongside aberrant PnC inhibitory neurotransmission, are associated with the PPI deficits observed in *Prodh^−/−^*mice.

## Discussion

Overall, we used the Cal-Light technique for the first time in the mouse CeA to photo-tag, quantify, chemically characterize, and manipulate the activity of neurons specifically activated during PPI, with high spatiotemporal precision. Our *in situ* hybridization results identified the glutamatergic nature of PPI-related CeA neurons, confirming that these CeA glutamatergic neurons must activate a post-synaptic inhibitory mechanism to contribute to PPI. We also targeted PnC neurons with Cal-light and discovered a greater proportion of active PnC inhibitory neurons during PPI (possibly recruited by CeA inputs) compared to baseline startle. Given that amygdala dysfunctions are common in diseases with PPI deficits, we hypothesized that CeA-PnC synapses could be affected in such diseases. To test this hypothesis, we used the *Prodh^−/−^* mouse, a schizophrenia-relevant disease model that exhibits amygdala impairments and PPI deficits. Surprisingly, by optogenetically activating CeA-PnC glutamatergic synapses, we were able to rescue PPI deficits in *Prodh^−/−^* mice. Finally, we demonstrated that in *Prodh^−/−^* mice, PnC-projecting CeA neurons exhibit increased axonal branching and excitability. Taken together, these *ex vivo* cytoarchitectural, histochemical, *in vivo* behavioral, and *in vitro* electrophysiological results provide, for the first time, insights into the mechanism by which the schizophrenia*-*relevant *Prodh* deficiency alters the structure and function of CeA-PnC glutamatergic synapses and impacts PPI.

### The contribution of the CeA to acoustic startle and PPI

The medial CeA (CeM) serves as the primary output center of the amygdala, transmitting integrated information to various premotor hindbrain regions that drive both innate and learned defensive behaviors (Swanson and Petrovich, 1998; Isosaka et al., 2015; Tillman et al., 2018). Previous research in rats has shown that neurons in the CeM can ***enhance*** the acoustic startle response by sending inputs to startle-mediating PnC giant neurons (Rosen et al., 1991; Koch and Ebert, 1993). In contrast, using tract-tracing and *in vivo* optogenetics in mice, we recently demonstrated that medial CamKIIα^+^ CeA neurons send direct glutamatergic projections to PnC glycinergic neurons and can ***decrease*** a subsequent startle response and thereby contribute to PPI (Cano et al., 2021; Huang et al., 2024). Consistent with these findings, we used Cal-Light, a calcium-dependent and blue light-sensitive method, to specifically identify and manipulate CeA neurons active during PPI and confirmed their glutamatergic nature via *in situ* hybridization. Fung et al. (2011) also reported CeA neurons sending glutamatergic projections to pontine reticular nucleus in guinea pigs, indicating a role of CeA glutamatergic neurons in regulating brainstem functions. While the CeA also contains a large population of GABAergic neurons that send inputs to various brainstem regions (Swanson and Petrovich, 1998; Saha et al., 2000; Jüngling et al., 2015; McCullough et al., 2018), no CeA GABAergic projections have been found to extend into the PnC, further highlighting the importance of the CeA-PnC glutamatergic neurons in PPI.

Aside from labeling active PnC-projecting CeA neurons, we were also able to functionally assess the contribution of these neurons, as the Cal-Light construct also contains the Halorhodopsin (NpHR 3.0) reporter, a yellow light-sensitive inhibitory optogenetic tool. Our results show that the photo-inhibition of Cal-Light/ NpHR 3.0^+^ CeA neurons reduced PPI without affecting the startle response itself. These findings not only confirm the involvement of CeA glutamatergic neurons in PPI but also demonstrate that Cal-Light enables the investigation of neural circuits underlying short-lived, transient behaviors (Lee et al., 2017; Hyun et al., 2022). Additional experiments are needed to identify the upstream pathways and brain structures that activate PPI-related CeA neurons.

### PnC neurons active during acoustic startle and PPI

We also utilized the Cal-Light technique to label and visualize PnC neurons specifically linked to acoustic baseline startle and PPI. While the PnC includes a heterogeneous population of small and medium-sized neurons (soma size between 20-30 µm in diameter; Zeilhofer et al., 2005), previous electrophysiological recordings focused solely on PnC giant glutamatergic neurons, identified by their larger soma size exceeding 30 µm (size range: 32-83 µm; mean = 44 µm; Lingenhöhl and Friauf, 1992; Schmid et al., 2010; Geis and Schmid, 2011; Zaman et al., 2017; Zheng and Schmid, 2023). These studies have provided essential insights into the function of PnC giant neurons during startle responses. However, understanding how these giant neurons, which represent about 1% of the PnC neuronal population (Davis et al., 1982; Lingenhöhl and Friauf, 1992; Koch and Ebert, 1993; Lee et al., 1996; Li et al., 2009), are inhibited during PPI has been limited by the lack of investigation into the contributions of non-giant PnC neurons, including inhibitory neurons, within the startle circuit. Here, for the first time, our results revealed that in addition to large (and likely startle-mediating) PnC giant neurons, 15% of Cal-Light-targeted PnC neurons active during acoustic baseline startle are inhibitory and PV^+^. Since we previously showed that GlyT2^+^ PnC inhibitory neurons do not contribute to baseline startle (Cano et al., 2021; Huang et al., 2024) and that PV^+^ neurons include both GlyT2^+^ and GABAergic inhibitory neurons, future studies should chemically identify the nature of the PV^+^ PnC inhibitory neurons active during baseline startle.

To further explore the inhibitory mechanisms underlying PPI, we also used Cal-Light to identify the population of PV^+^ PnC neurons active during PPI. Our results show that during PPI, the proportion of active PV^+^ PnC neurons rose to 25%, representing a 66% increase compared to the number of PV^+^ neurons active during startle alone. These additional PV^+^ neurons could include GlyT2^+^ PnC neurons, previously shown to contribute to PPI (Cano et al., 2021; Huang et al., 2024).

Based on these Cal-Light results, it is tempting to speculate that PV^+^/GlyT2^−^ PnC neurons contribute to startle responses, while PV^+^/GlyT2^+^ neurons underlie PPI. This would suggest that specific subtypes of PV^+^ inhibitory neurons contribute to distinct behavioral functions and different behavior-dependent network activities (Freund and Katona, 2007). Indeed, previous studies have examined how two genetically defined, partially overlapping but distinct populations of GABAergic interneurons, GAD65^+^ (expressing the GABA-synthesizing enzyme) and PV^+^ inhibitory neurons, affect different behavioral deficits relevant to schizophrenia. Their results showed that photo-inhibiting PV^+^/GAD 65^−^ inhibitory hippocampal neurons significantly reduced PPI, whereas photo-inhibiting PV^+^/GAD 65^+^ inhibitory neurons increased locomotor activity without affecting PPI (Nguyen et al., 2014).

### The schizophrenia-related *Prodh* dysfunction and PPI deficits

PPI deficits are a hallmark of schizophrenia. The proper functioning of the amygdala and glutamate neurotransmission is essential for PPI (Javanbakht, 2006; Cardno and Owen, 2014; Mena et al., 2016; Cano et al., 2021; Huang et al., 2024). However, the amygdala decreases in volume during the development of schizophrenia (Ellison-Wright et al., 2008; Meyer-Lindenberg, 2010). Furthermore, changes in amygdala activity and its functional connectivity with other regions, such as the medial prefrontal cortex, are linked to positive symptoms, disturbed social cognition, and impaired emotional regulation in schizophrenia (Lennox et al., 2000; Brunet-Gouet and Decety, 2006; Rasetti et al., 2009; Goghari et al., 2010; Li et al., 2010; Gruber et al., 2014). Aside from glutamate, both human and animal studies report that schizophrenia dysregulates GABAergic inhibitory neurotransmission in the amygdala (Gilabert-Juan et al., 2011, 2012; Egbujo et al., 2016; Castillo-Gómez et al., 2017). While very few studies have explored the contribution of glycinergic neurotransmission to schizophrenia, systematic glycine administration has consistently been shown to improve both positive and negative symptoms associated with the disorder (Heresco-Levy et al., 1999; Singh and Singh, 2011; Harvey and Yee, 2013).

The 22q11.2 deletion syndrome (22q11.2DS), caused by a microdeletion on chromosome 22, is one of the most significant known risk factors for schizophrenia. Individuals with 22q11.2DS have a 20-25 times higher likelihood of developing schizophrenia compared to the general population (Bassett et al., 2000; Hodgkinson et al., 2001). In fact, approximately one-third of people with 22q11.2DS develop schizophrenia or schizoaffective disorder, representing about 1-2% of all schizophrenia cases (Bassett and Chow, 2008; Karayiorgou et al., 2010). Located within the 22q11.2 deletion region, the *PRODH* gene has been identified as a susceptibility gene for schizophrenia (Kempf et al., 2008; Roussos et al., 2009; Ghasemvand et al., 2015). It encodes proline dehydrogenase, the rate-limiting enzyme in proline degradation. In humans, a deficiency in *PRODH* leads to an accumulation of proline in the blood and brain (Kempf et al., 2008; Roussos et al., 2009; Ghasemvand et al., 2015). Similarly, *Prodh^−/−^* mice show increased proline levels in the hypothalamus, hippocampus, frontal cortex, and amygdala (Gogos et al., 1999). *Prodh* deficiency reduces glutamatergic neurotransmission in the hippocampus (Paterlini et al., 2005) and GABAergic neurotransmission in the medial prefrontal cortex (Crabtree et al., 2016). Reminiscent of schizophrenia patients, *Prodh^−/−^* mice also exhibit impaired fear conditioning and reduced PPI levels, both of which rely on proper amygdala functioning (Gogos et al., 1999; Paterlini et al., 2005; Hall et al., 2009).

Previous findings of structural alterations were also associated with behavioral impairments and changes in electrophysiological recordings at cortical synapses of *Df(16)A^+/−^* mice, a model of the 22q11.2 microdeletion (Fénelon et al., 2011, 2013; Mukai et al., 2008, 2015). Our immunohistochemical analysis in *Prodh^−/−^* mice shows an increase in the CeA axonal branching pattern (which is disrupted by various genes within the 22q11.2 locus). Associated with these structural changes, our whole-cell patch-clamp recordings demonstrated increased intrinsic excitability of CeA glutamatergic neurons projecting to the PnC in *Prodh^−/−^* mice. However, *Prodh^−/−^* mice exhibit reduced PPI, implicating decreased inhibitory neurotransmission in the PnC. One possibility is that the excitatory CeA glutamatergic inputs are enhanced and activate additional PnC inhibitory mechanisms (including GlyT2^+^ neurons), to compensate for insufficient inhibitory neurotransmission in the PnC. Another possibility is that in addition to recruiting GlyT2^+^ neurons, the abnormal excitability and axonal branching of CeA neurons aberrantly recruit PnC excitatory neurons, leading to a reduction in PPI. Overall, while further studies should test these possibilities, our findings offer mechanistic insights into how *Prodh* deficiency alters the CeA-PnC glutamatergic connection and affects PnC inhibitory networks contributing to PPI deficits.

## MATERIALS and METHODS

### Animals

Experiments were performed on C57BL/6 mice (N = 52; The Jackson Laboratory, Bar Harbor, ME), GlyT2-eGFP mice (N = 5; graciously provided by Dr. Manuel Miranda-Arango, University of Texas at El Paso, El Paso, TX) and *Prodh^−/−^*mice (N = 23; graciously provided by Dr. Joseph A. Gogos, Columbia University, New York). Litters were weaned at PND 21 and housed together until stereotaxic surgeries were performed. Mice received food and water ad libitum in a 12-hour light/dark cycle from 8:00 am to 8:00 pm. All the stereotaxic coordinates and cytoarchitectural boundaries were derived from Paxinos and Franklin Mouse Brain Atlas. After surgical procedures, the mice were individually housed and monitored throughout the recovery period. All experimental protocols were conducted in accordance with and approved by the Institutional Animal Care and Use Committee of the University of Massachusetts Amherst.

### Surgical procedure and viral constructs

Mice were sedated using 5% isoflurane vapors (Piramal Critical Care, Bethlehem, PA) and then placed on a stereotaxic apparatus (model 900, David Kopf, Tujunga, CA) with ear bars and a nose cone for immobilization. Throughout the surgical procedure, mice were maintained under 1.5%-2.5% isoflurane. The head of the mice was leveled on all three axes relative to bregma. A craniotomy was performed directly dorsal to the injection site, and viral vectors were delivered using a microinjector (Stoelting Co., Wood Lane, IL) with a 5μl Hamilton syringe (Hamilton Company Inc., Reno, NV) and a 32-gauge steel needle.

To transduce CeA (AP −1.35 mm, ML +2.66 mm, DV +4.6 mm) neurons with Cal-Light components, 500 nl of Cal-Light viral vectors (AAV1-hSYN-TM-CaM-TEV-N-AsLOV2-TEVseq-tTA : AAV1-hSYNM13-TEV-C-P2A-tdTomato : AAV1-TRE-eNpHR-eGFP = 1 : 2 : 2; Vigene Biosciences) were injected unilaterally in the CeA of C57BL/6 mice (N = 5). To transduce PnC (AP −5.35 mm; ML +0.5 mm, DV +5.15 mm) neurons with Cal-Light viral components, 750 nl of Cal-Light viral vectors (AAV1-hSYN-TM-CaM-TEV-N-AsLOV2-TEVseq-tTA : AAV1-hSYN-M13-TEV-C-P2A-tdTomato : AAV1-TRE-eNpHR-eGFP = 1 : 2 : 2; Vigene Biosciences) were injected unilaterally in the PnC of C57BL/6 mice (N = 23).

For optogenetic stimulation of CeA glutamatergic neurons, C57BL/6 mice (N = 6) or *Prodh^−/−^* mice (N = 9) unilaterally injected with 100-120 nl of the AAV-CaMKIIα-hChR2(H134R)-mCherry (Addgene, catalog #26975) viral vector in the CeA (AP −1.35 mm, ML +2.66 mm, DV +4.6 mm). In addition, to retrogradely label PnC-projecting CeA glutamatergic neurons for *in vitro* patch clamp recordings, 100-120 nl of rgAAV-CaMKIIα-hChR2(H134R)-mCherry (Addgene, catalog #26975) viral vectors were injected unilaterally in the PnC (AP −5.35 mm; ML +0.5 mm, DV +5.15 mm) of C57BL/6 mice (N = 7) or *Prodh^−/−^* mice (N = 8). After viral injection, the microinjection syringe was left in place for 10 mins after infusion to limit spillover during needle retraction.

Three weeks after the viral injection, mice were sedated by inhaling 5% isoflurane vapors, placed in a stereotaxic apparatus, and immobilized using ear bars and a nose cone. Mice were maintained under anesthesia (1.5%-2% isoflurane), and the head was leveled in all three axes with bregma as a reference. A craniotomy was drilled directly dorsal to the implantation site, and a cannula guide with a 200 μm core optical fiber (Thorlabs, Newton, NJ) was implanted at the level of the PnC (AP −5.35 mm, ML +0.5 mm, DV +4.75 mm) or at the level of the CeA (AP −1.35 mm, ML +2.66 mm, DV +4.2 mm). The cannula guide was cemented to the skull with dental cement (Parkell, Edgewood, NY). Mice were allowed to recover for 7 days post-surgery before behavioral testing.

### Behavioral measurement

The measurement of the acoustic startle response (ASR) and PPI was conducted in an enclosed sound-attenuated startle chamber from PanLab (Harvard Apparatus, Holliston, MA). Sound pressure levels were calibrated using a standard SPL meter (model 407730, Extech, Nashua, NH). Mice were placed on a movement-sensitive platform, and vertical displacements of the platform induced by startle responses were converted into a voltage trace by a piezoelectric transducer located underneath the platform. Startle amplitude was measured as the peak-to-peak maximum startle magnitude of the signal recorded during a 1s window following the presentation of the acoustic stimulation. All the testing trials were designed and recorded using PACKWIN V2.0 software in PanLab System (Harvard Apparatus, Holliston, MA). Prior to any testing session, mice were handled and acclimatized to the testing chamber, where they were exposed to a 65dB background noise for 10 min.

### Testing of the acoustic startle response and prepulse inhibition

During the ASR task, mice were presented with 40 ms sound pulses at different intensities (in dB: 70, 80, 90, 100, 110, and 120) every 15 s, in a pseudorandomized order. Background noise (65 dB) was presented during the 15 s intervals between sounds. For each task, a total of 30 trials (i.e., 6 intensities, each presented 5 times) were acquired and quantified. During the PPI task, non-startling prepulses (75 dB noise; 20 ms) followed by startling pulses (120 dB noise; 40 ms) were presented at various inter-stimulus intervals (ISIs, in ms: 10, 30, 50, 100, 200, 300, and 500) every 29 s, in a pseudorandomized order. Background noise (65 dB) was presented during the 29 s intervals between sounds. For each task, a total of 49 trials (i.e., 7 ISIs, each presented 7 times) were acquired and quantified. Additionally, baseline startle amplitude was measured with only the startling pulses being presented.

### Photo-tagging of active neurons expressing Cal-Light

During the ASR task used for the optical labeling of active neurons, a series of 30 startling pulses at 120 dB (40 ms) were presented at regular 15 s intervals, accompanied by a background noise of 65dB during the intervals between sounds. To photo-label PnC neurons active during ASR, constant 473 nm blue light (Opto Engine LLC, Midvale, UT) was delivered during the 40 ms startling pulse (120 dB). Mice completed the ASR task paired with blue light 5 times per day for 5 consecutive days.

During the PPI task used for the optical labeling of active neurons, a combination of non-startling prepulses (75 dB noise; 20 ms) followed by startling pulses (120 dB noise; 40 ms) were presented at different ISIs (i.e., 30, 50, and 100 ms) known to effectively elicit measurable PPI (based on studies by Valsamis and Schmid, 2011; Azzopardi et al., 2018; Cano et al., 2021; Huang et al., 2024). The presentation of these stimuli occurred every 29 s in a pseudorandomized order, accompanied by a background noise of 65 dB during the intervals between sounds. For each task, a total of 30 trials (i.e., 3 ISIs, each presented 10 times) were acquired and quantified. Additionally, baseline startle amplitude was measured with only the startling pulses being presented. To photo-label CeA neurons active during PPI, constant 473 nm blue light (Opto Engine LLC, Midvale, UT) was delivered during the 20 ms non-startling prepulse (75 dB). To photo-label PnC neurons active during PPI, constant 473 nm blue light (Opto Engine LLC, Midvale, UT) was delivered concurrently with the 20 ms non-startling prepulse (75 dB), lasting throughout the entire ISI. Mice completed the PPI task paired with blue light 3 times per day for 5 consecutive days, followed by a 2-day resting period before the next behavioral testing (optical inhibition).

### Photo-inhibition of active neurons expressing Cal-Light

To photo-inhibit labeled neurons during acoustic baseline startle responses, a train of yellow light pulses (5 Hz, 3 ms ON, 197 ms OFF; yellow 593.5 nm laser, Opto Engine LLC, Midvale, UT) was delivered 400 ms before and covered the entire acoustic startling pulse. The ASR task was conducted on 2 consecutive days, both in the absence and presence of yellow light. Then, to photo-inhibit labeled neurons during PPI, a train of yellow light pulses (5 Hz, 3 ms ON, 197 ms OFF; yellow 593.5 nm laser, Opto Engine LLC, Midvale, UT) was delivered 400 ms before the acoustic non-startling prepulse, lasting the entire ISI. The PPI task was conducted on 2 consecutive days, both in the absence and presence of yellow light.

### Photo-activation of CeA-PnC glutamatergic synapses

To photo-activate the CeA-PnC glutamatergic connection, a train of blue light pulses (5 Hz, 3 ms ON, 197 ms OFF; blue 473 nm laser, Opto Engine LLC, Midvale, UT) was delivered 400 ms before the acoustic non-startling prepulse, lasting the entire ISI. The PPI levels were measured both in the absence of and in the presence of blue light. During the PPI task using optical stimulation as prepulses, a train of blue light pulses (3 pulses, 15 ms ON, 5 ms OFF; blue 473 nm laser, Opto Engine LLC, Midvale, UT) was delivered at 50 Hz before the startling pulse. The ISIs between the light prepulse and the startling pulse were (from the end of the prepulse to the onset of the pulse; in ms): 10, 30, 50, 100, 200, 300, and 500.

At the end of each experiment, histological analyses were performed to confirm that: 1) the confined localization of the injected viral vectors within the region of interest, and 2) the cannula guide placement was successfully aimed at the region of interest. If these criteria were not met, the subject was excluded from the study.

### Immunohistochemistry

Mice were perfused transcardially with 0.9% saline solution for 10 min, followed by 4% paraformaldehyde (PFA) in 0.1 M phosphate buffer saline (PBS; pH 7.4) for 15 min. The brains were extracted and post-fixed overnight in 4% PFA solution, then embedded in 30% sucrose in PBS for 2 days at 4 °C. Subsequently, the brains were frozen in chilled hexanes for 1 min and stored at −80 °C. Coronal brain sections with a thickness of 30 μm were obtained using a microtome (Leica CM3050 cryostat) and stored in cryoprotectant solution (50% 0.05 M phosphate buffer, 30% ethylene glycol, 20% glycerol) at −20 °C.

To stain for PRODH, coronal tissue sections were washed with 0.1 M Tris-buffered saline (TBS; pH 7.4) and then incubated in blocking solution (2% normal donkey serum, 0.1% Triton X-100; in 0.1 M TBS) for 2 h at room temperature. The tissue sections were then incubated with a goat anti-PRODH primary antibody (1:1000, Sigma Aldrich, catalog #SAB2501795-100UG) for 72 h at 4 °C, washed with TBS, and subsequently incubated in a Cy3-conjugated donkey anti-goat secondary antibody (1:500, Jackson ImmunoResearch Laboratories, catalog #706165147) for 4 h at room temperature. To stain for parvalbumin, tissue sections were incubated with a rabbit anti-parvalbumin primary antibody (1:1000, Abcam, catalog #ab11427), followed by incubation with a Cy5-conjugated donkey anti-rabbit secondary antibody (1:500, Jackson ImmunoResearch Laboratories, catalog #705175147). In the end, brain slices were rinsed with TBS, mounted, and coverslipped to visualize injection and projection sites.

### *In situ* hybridization

Mice injected with Cal-Light viral vectors in the CeA (N = 5) were perfused and coronal sections (30 μm) at the level of the CeA were cut using a microtome and stored in cryoprotectant solution at −20 °C as described above. Next, tissue sections were washed with TBST (0.5x TBS + 0.1% Tween 20) once and mounted onto superfrost plus slides. Then slides were dry at room temperature for 1 h, submerged into ddH_2_O for 3 times, dry at room temperature for 1 h and baked at 60 °C in a circulating air oven for another hour. Next, slides were submerged into freshly prepared cold 4% PFA for 1 h and dehydrated in increasing ethanol solutions (50%, 70%, 100%, 100%; 5 min each at room temperature). Then, the RNAscope assay (Advanced Cell Diagnostics) started by incubating in hydrogen peroxide (H_2_O_2_) for 10 min in a humidified chamber, followed by protease III incubation for 20 min. RNA hybridization probes against genes encoding mouse VGLUT2 (1170921-C1), eYFP (312131-C2), and Tdtomato (317041-C3) were then incubated for 2 h at 40 °C. Antisense probes were also included as controls in a separate glass slide. Probe signals were then developed separately with TSA Vivid Dyes (TSA 650 1:1000, TSA 520 1:1500, TSA 570 1:1500) and coverslipped with ProLong Gold^TM^ with DAPI.

### Imaging analysis

Imaging was conducted using an upright confocal laser-scanning microscope (A1R-TIRF, Nikon) equipped with 405, 445, 488, 514, 561, 640 nm lasers and 10x, 20x, and 40x objectives. The cell counting analysis was performed using ImageJ (NIH). Gaussian blur filters, thresholding, watershed and denoise function were applied on images. The processed images were then measured by particle analyzer. Image calculator with appropriate operations was used to process images for measurements of double-positive signals. A total of 1224 images (both fluorescence and bright-field) were analyzed for each brain region, and boundaries were delineated using Adobe Illustrator (Adobe, San Jose, CA).

### Electrophysiological recordings

Experiments were performed on adult *Prodh^−/−^* mice and their WT littermates 3-5 weeks after rgAAV-CaMKIIα-hChR2(H134R)-mCherry injection in the PnC. The mice were anesthetized, followed by cervical dislocation and rapid decapitation. Then, the brain was harvested and placed in ice-cold dissecting solution with the following composition (in mM): sucrose (195), NaCl (10), KCl (2.5), NaH_2_PO_4_ (1), NaHCO_3_ (25), glucose (10), MgSO_4_ (4), CaCl_2_ (0.5), constantly bubbled with 5% CO_2_/95% O_2_. Then, 300 μm thick coronal sections containing the CeA were cut using a vibratome (Leica VT 1200S, Wetzlar, Germany). These CeA sections were immediately transferred to a beaker filled with artificial cerebrospinal fluid (aCSF) at room temperature bubbled with 5% CO_2_/95% O_2_ with the following composition (in mM): NaCl (124), KCl (2.5), NaH_2_PO_4_ (1), NaHCO_3_ (25), glucose (10), MgSO_4_ (1), CaCl_2_ (2). Slices were transferred to an electrophysiological recording setup and allowed to recover on an interface chamber for at least 2 h at 31-32 °C. aCSF was fed by gravity at a rate of ≈ 2 mL/min.

<u>Whole-cell recordings</u> (N = 8 *Prodh^−/−^* mice, n = 12 CeA neurons; N = 7 WT mice, n = 12 CeA neurons) were performed using glass pipettes (3-5 MΩ) filled with intracellular solution (in mM): KMeSO4 (125), KCl (10), HEPES (10), NaCl (4), EGTA (0.1), MgATP (4), Na_2_GTP (0.3), Phosphocreatine (10), Biocytin (0.1%) (pH = 7.3; osmolarity = 285-300 mosm). The glass micro-electrode was mounted on a patch clamp headstage (Molecular Devices LLC, Sunnyvale, CA; catalog# CV-7B), which was attached to a multi-micromanipulator (Sutter Instrument, Novato, CA; catalog# MPC-200). Data were acquired with pClamp10 software using a MultiClamp™ 700B amplifier (Molecular Devices LLC, Sunnyvale, CA) and a Digidata 1550B digitizer (Molecular Devices LLC, Sunnyvale, CA). CaMKIIα-ChR2-mCherry^+^ CeA cells were imaged and targeted using NIS-Elements Basic Research software (version 5.11, Nikon Instruments Inc., Melville, NY). Only cells with an initial seal resistance greater than 1 GΩ, a resting membrane potential between −60 mV to −70 mV, and a holding current within −100 pA to 100 pA at resting membrane potential and overshooting action potentials were used.

In CeA slices, intrinsic properties were recorded after a 10-min equilibration period. Then, 15 pA depolarizing current steps were injected for 500 ms to induce action potentials in CeA neurons expressing CaMKIIα-ChR2-mCherry in the current clamp. Spontaneous EPSCs were recorded at a holding potential of −70 mV, in the voltage clamp. Evoked EPSPs were also recorded in these CeA neurons held at −70 mV, in response to a 1ms blue light pulse. Blue light was delivered every 30 s using a 200 μm optic fiber mounted on a micromanipulator connected to a blue LED (473 nm; Plexon, Dallas, TX) and positioned in close proximity to the recorded neuron.

At the end of all whole-cell recordings, the cell membrane was sealed by forming an outside-out patch. The glass microelectrode was slowly retracted, and as the series resistance increased, the membrane potential was clamped at −40 mV. The 300 μm thick acute brain slices containing the recorded cells were immersed in 4% PFA solution overnight. Following overnight PFA fixation, these brain slices were rinsed with PBS (3 times, 5 min each). Slices were then incubated in an anti-RFP antibody and in a complementary secondary antibody to enhance the fluorescence of the viral vectors used. Following PBS rinses, slices were incubated with Cy5-conjugated streptavidin (a biotin binding protein) diluted in PBS (with 0.1% Triton X-100) at room temperature for 4-5 h or overnight at 4 °C. Slices were then rinsed with PBS, mounted on glass slides, coverslipped, and sealed with ProLong^TM^ Gold antifade reagent (Invitrogen by Thermo Fisher Scientific, Waltham, MA, catalog# P36934), and airdried overnight in the dark.

### Statistical analysis

Statistical analyses were conducted using GraphPad Prism (GraphPad Software, La Jolla, CA). Normality and equal variance assumptions were tested, and if not met, equivalent nonparametric tests were used. For repeated-measures (RM) ANOVA, Mauchly’s test of sphericity was conducted, and if violated, Greenhouse-Geisser or Huynh-Feldt corrections were applied. *Post hoc* tests were performed using Student’s t-test with Bonferroni correction when necessary. The significance criterion was set at 0.05.

For the results from the *in vivo* startle response tasks, two-way RM ANOVA was conducted to assess the effects of sound intensity and optical manipulation on the startle amplitude. Mixed ANOVA was conducted to assess the effects of sound intensity on the startle amplitude between *Prodh*^−/−^ and WT mice. For the *in vivo* PPI tasks, two-way RM ANOVA was conducted to evaluate the effects of ISI and optical manipulation on PPI values. Mixed ANOVA was used to compare PPI levels measured with or without photo-activation of the CeA-PnC connection in *Prodh*^−/−^ with baseline PPI levels in WT mice. PPI was quantified as follows: %PPI = [(1 - startle amplitude during “Prepulse + Pulse” trials / startle amplitude during “Pulse only” trials) × 100], which represents the extent of startle suppression due to the presentation of the prepulse. Results from *in vitro* whole-cell patch clamp recordings under voltage clamp conditions were quantified by the amplitude and frequency of the maximum current peak. In current clamp, two-way ANOVA was conducted to assess the effects of optical manipulation on the spiking activity of light unresponsive and responsive cells. Differences in intrinsic and synaptic properties between *Prodh*^−/−^ and WT mice were evaluated with a paired t-test. Sample sizes were determined based on expected outcomes, variances, and power analysis. Data are presented as means ± SEM.

## Notes

### Competing Interest Statement

The authors have declared no competing interest.

### Funding

- Armstrong Funds for Science

