## Supplemental Information for "Targeting Amygdala-Brainstem Synapses to Reverse Prepulse Inhibition Deficits"

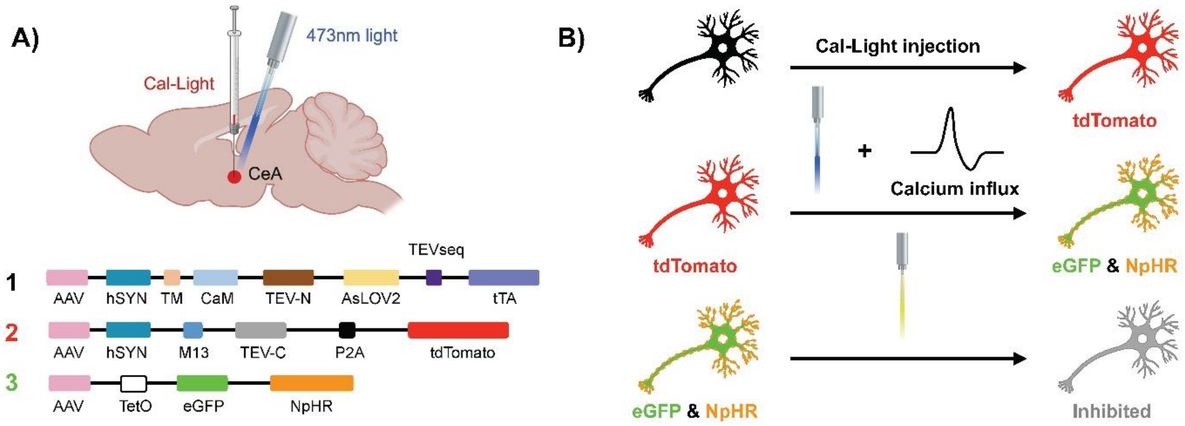


**Supplemental Figure 1. The Cal-Light system injected in the mouse brain.**

**(A)** Schematic illustration of the mouse brain sagittal section showing the injection of the Cal-Light virus and delivery of blue light. In the Cal-Light system, the TEV protease is split into two halves, with M13 and CaM proteins fused to its C- and N-termini, respectively. When $Ca^{2+}$ rises in the cytosol, M13 and CaM bind to each other and TEV protease regains proteolytic function. Upon blue light exposure, TEVseq is unmasked due to conformational change can thus be recognized by TEV protease. Then cleaved tTA translocates to the nucleus and binds to TetO, which initiates gene expression of eGFP and NpHR (Lee et al., 2017). **(B)** Schematic illustration showing the activation of Cal-Light system. ***Top,*** When Cal-Light is not activated, neurons only express tdTomato. ***Middle,*** When Cal-Light is activated in the presence of calcium and blue light, neurons will also express eGFP and NpHR. ***Bottom,*** Neurons expressing eGFP and NpHR can be photo-inhibited using yellow light.

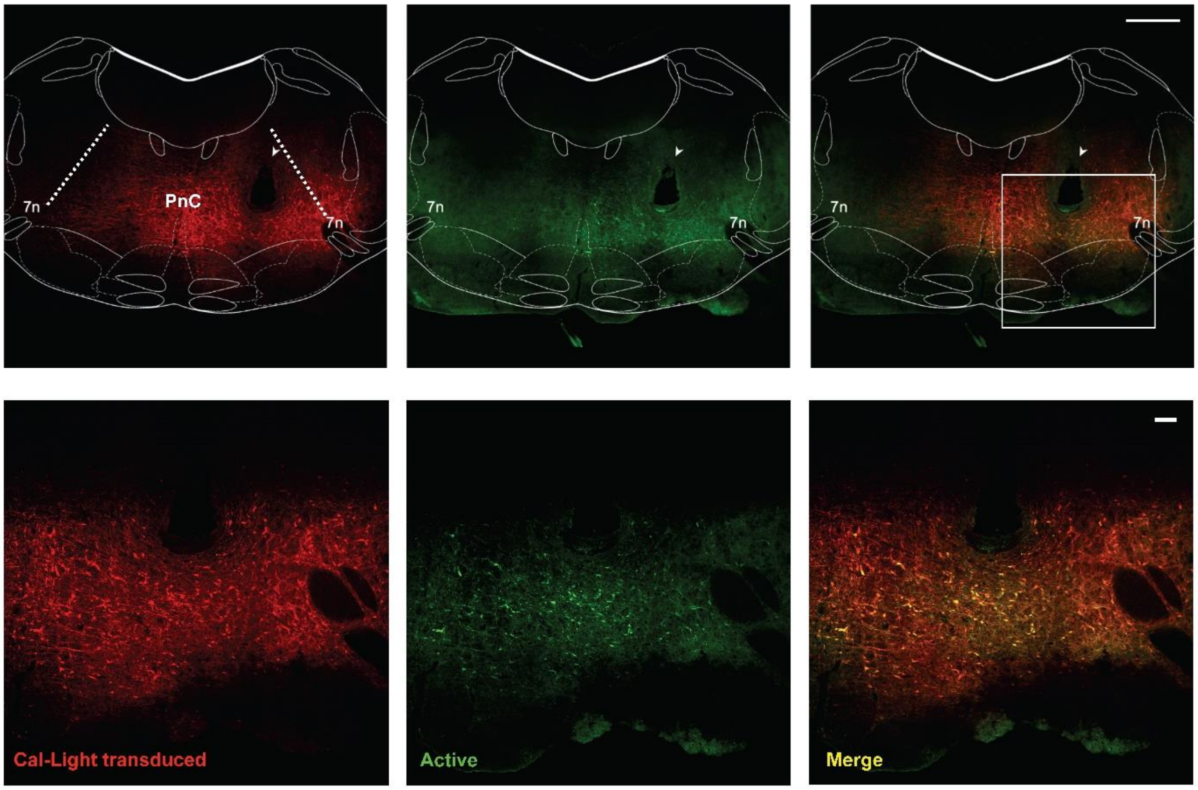


**Supplemental Figure 2. Identification of Cal-Light$^{+}$ PnC neurons active during baseline startle.**

***Top row***, In WT mice injected with Cal-Light in the PnC, only a subset of Cal-Light-targeted PnC neurons were active during the acoustic startle response (ASR). Images show PnC neurons active during ASR at low magnification. PnC neurons transduced with Cal-Light expressed tdTomato. In the presence of blue light, PnC neurons active during ASR expressed eGFP. The PnC is delineated by dashed lines. Arrowheads indicate the tract of implanted fiber optic for blue light delivery in the PnC. ***Bottom row***, Higher magnification images of the area in the white rectangle above. 7n: the 7$^{th}$ cranial nerve. Scale bars: 500µm (top row); 100µm (bottom row). Representative of N = 11 WT mice.

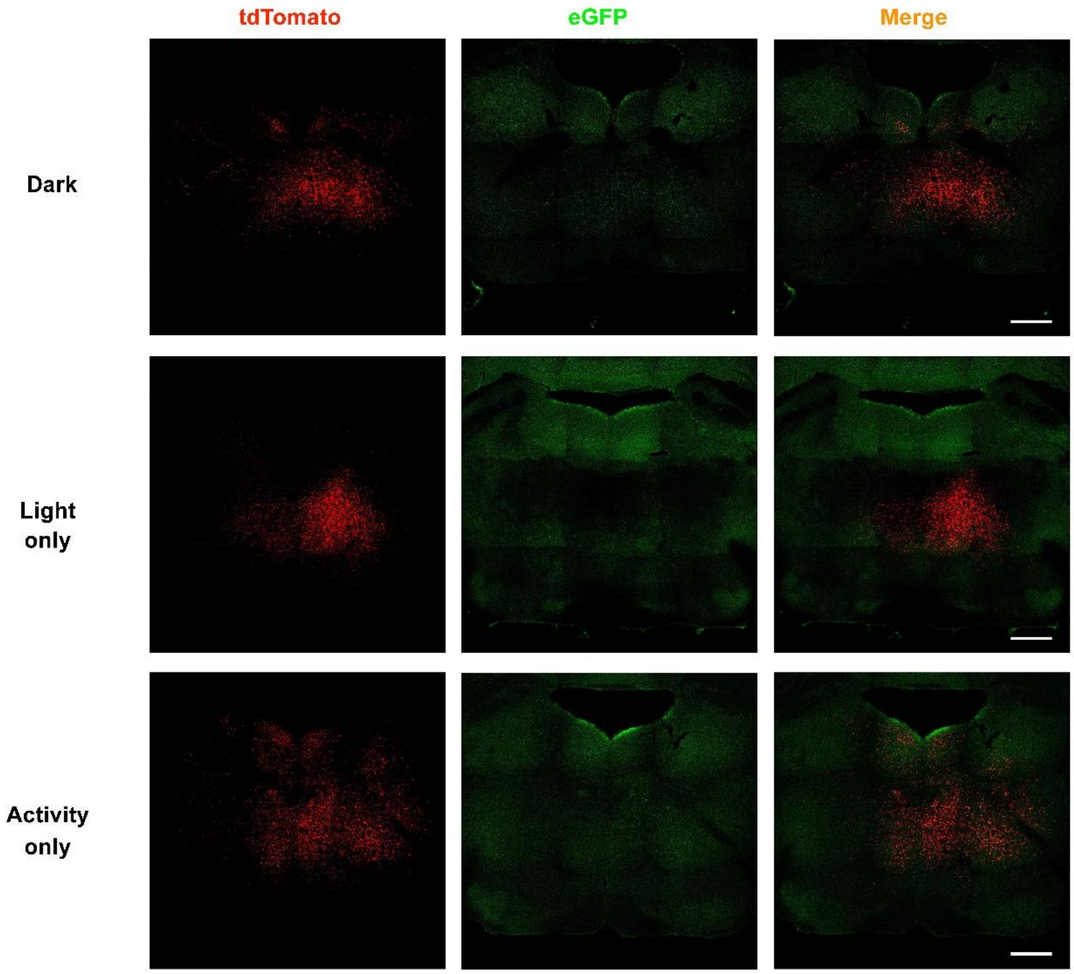


**Supplemental Figure 3. Cal-Light$^{+}$ neurons during control conditions**.

Representative images of Cal-Light$^{+}$ PnC neurons from mouse brains not exposed to blue light nor to acoustic stimulation (*top row*), from mouse brains exposed to blue light but not to acoustic stimulation (*middle row*), or from mouse brains not exposed to blue light during acoustic PPI. Scale bars: 500µm. Representative of N = 4 WT mice per condition.

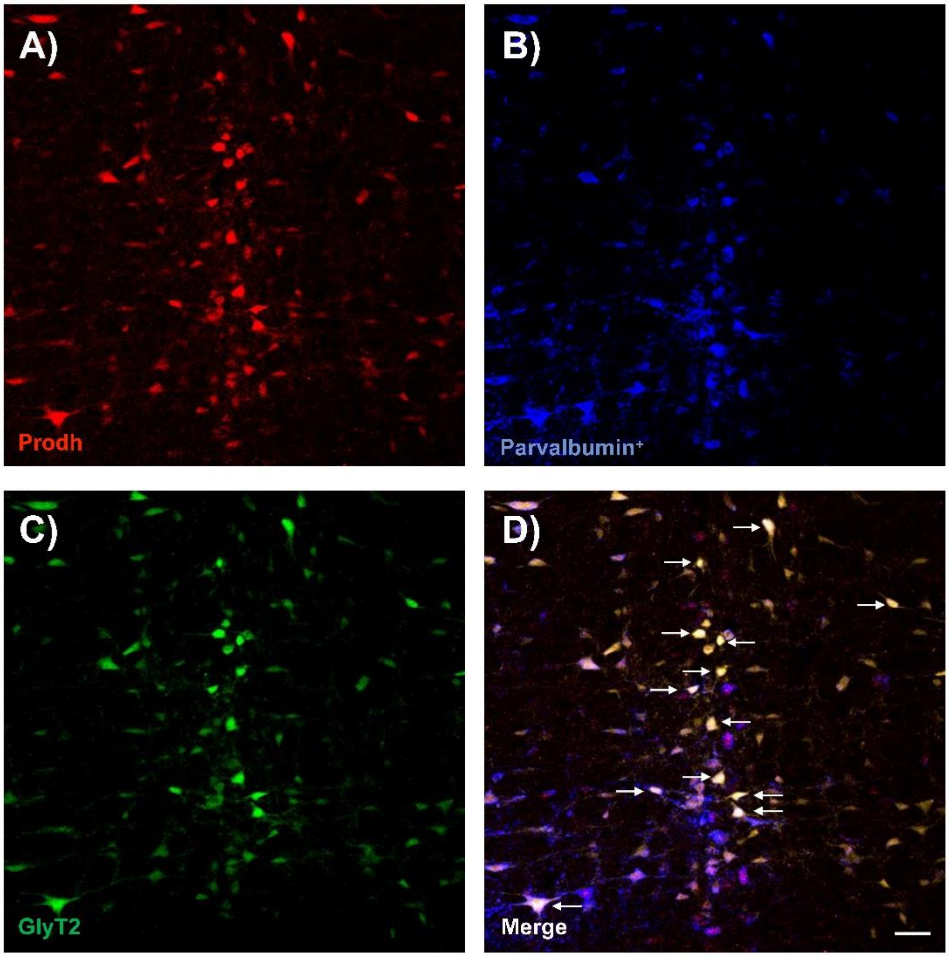


**Supplemental Figure 4.** ***Prodh*** **co-localizes with the inhibitory marker parvalbumin (PV) and GlyT2-eYFP in PnC neurons.**

**A-C)** Representative images showing co-labeling of **A)** *Prodh*, **B)** Parvalbumin and **C)** GlyT2-eYFP in the PnC of a GlyT2$^{Cre}$ mouse. **D)** Merged image with arrows identifying triple-labeled PnC neurons. Representative of N = 5 GlyT2$^{Cre}$ mice. Scale bar = 50µm**.**

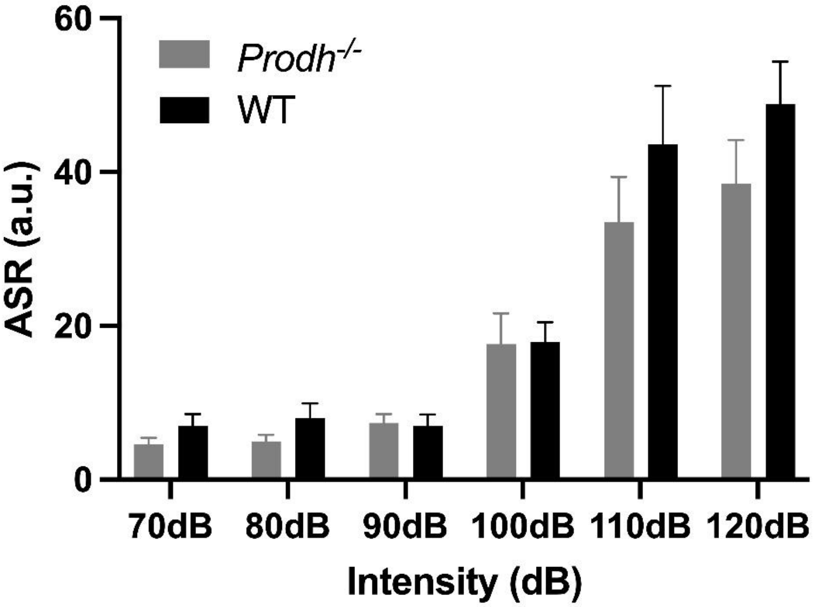


**Supplemental Figure 5**. ***Prodh* deficiency does not affect baseline startle.**

Graph showing that mean startle amplitude as a function of sound intensity does not differ between *Prodh*$^{-/-}$ (grey bars) and WT (black bars) mice [genotype: $F_{(1,13)} = 1.564$, $p = 0.233$; sound intensity: $F_{(5,65)} = 46.8$, $p < 0.001$; genotype x sound intensity interaction: $F_{(5,65)} = 0.91$, $p = 0.48$; Mixed ANOVA]. N = 9 *Prodh*$^{-/-}$ mice and N = 6 WT mice. Data are represented as mean ± SEM. $^*p < 0.05$, $^{**}p < 0.01$.

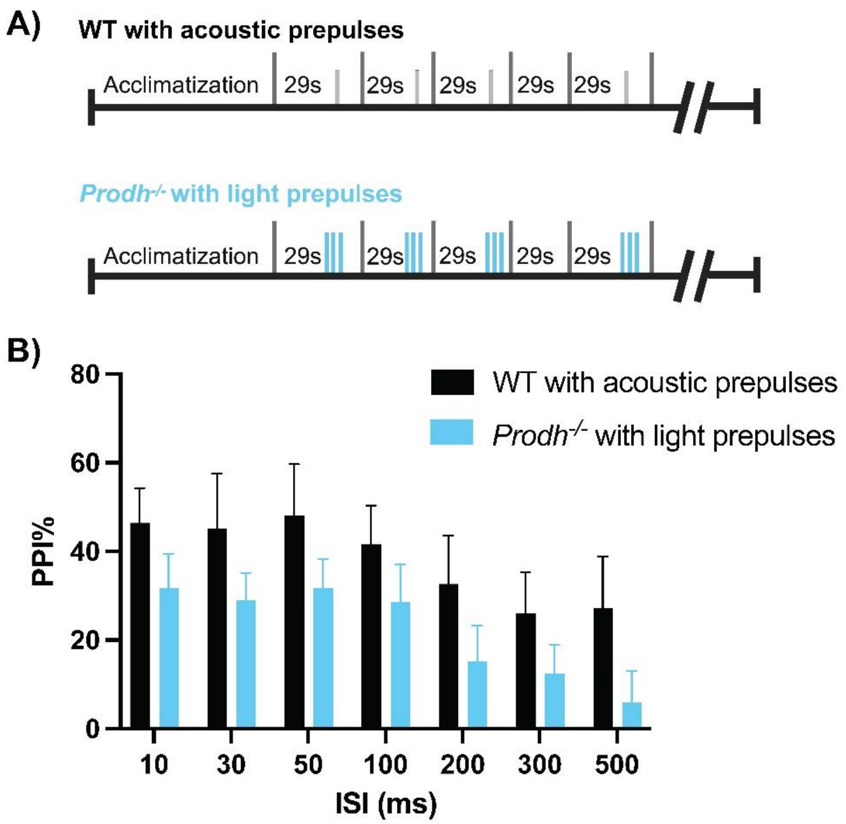


**Supplemental Figure 6**. **Photo-activation of CeA-PnC glutamatergic synapses is sufficient to induce WT-like acoustic PPI in *Prodh$^{-/-}$* mice.**

**A)** Schematic of the PPI protocols performed using acoustic prepulses in WT mice (***top***) or blue light prepulses in *Prodh$^{-/-}$* mice (**bottom**) injected with the blue-light sensitive excitatory optogenetic AAV-DJ-CamKIIα-hChR2-mCherry virus in the CeA. **B)** Graph showing mean PPI values as a function of ISIs, by using either sound in WT mice (black bars; "Acoustic prepulse") or blue-light photo-activation of CeA-PnC glutamatergic synapses as a prepulse in *Prodh$^{-/-}$* mice (blue bars; "Light prepulse"). The light-induced PPI in *Prodh$^{-/-}$* mice was comparable to the sound-induced PPI using acoustic prepulses in WT mice at all ISIs tested [Prepulse: $F_{(1,13)} = 2.519$, $p = 0.137$; ISI: $F_{(6,78)} = 6.974$, $p < 0.001$; Prepulse x ISI interaction: $F_{(6,78)} = 0.1373$, $p = 0.991$; Mixed ANOVA]. N = 9 *Prodh$^{-/-}$* mice and N = 6 WT mice. Data are represented as mean ± SEM. *$p < 0.05$.

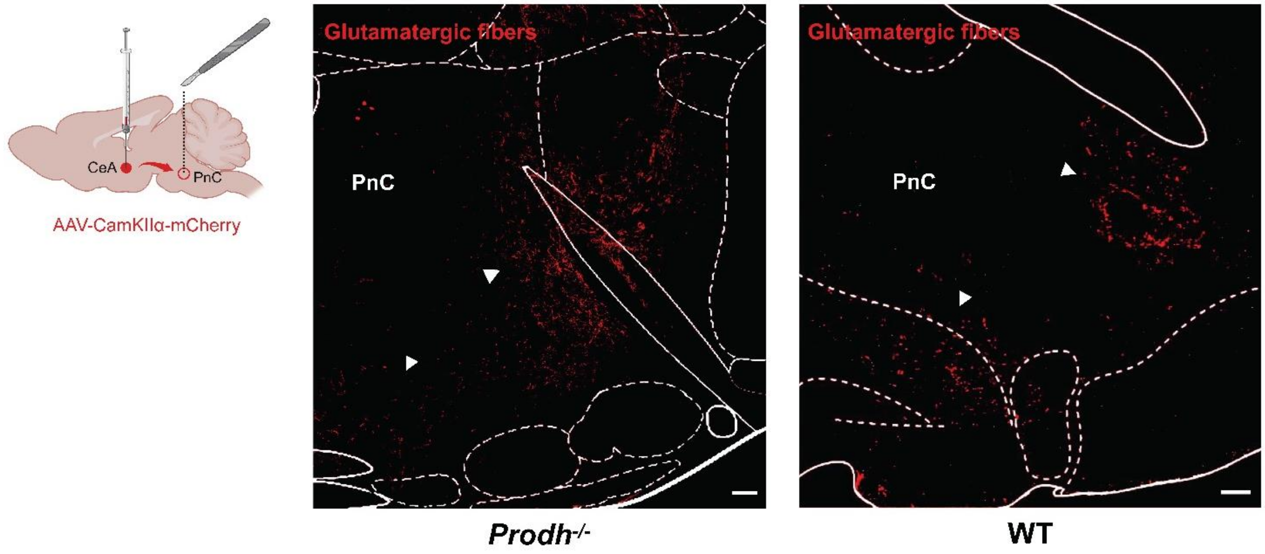


**Supplemental Figure 7**. **Increased number of CeA-PnC glutamatergic fibers in *Prodh$^{-/-}$* mice.**

Representative images showing the increased number of CeA glutamatergic fibers coursing into the ventrolateral PnC region of *Prodh$^{-/-}$* (***left***) compared to WT (***right***) mice. CeA glutamatergic neurons were transduced by a neuronal tracer that expresses mCherry under the control of the CamKIIα promoter. The white arrowheads indicate the location of CeA fibers coursing ventrally and laterally in the PnC. Scale bars: 100µm. Representative of 6 WT mice and 9 *Prodh$^{-/-}$* mice.

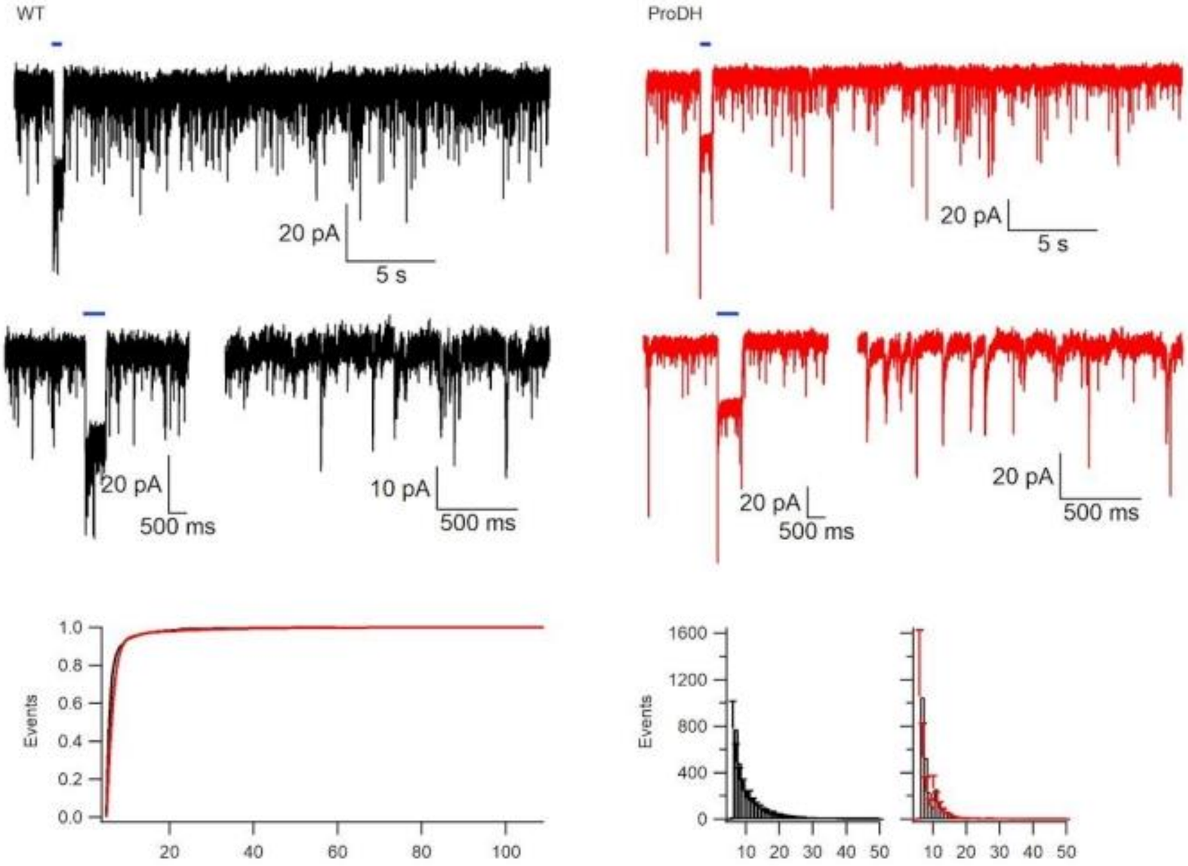


**Supplemental Figure 8. *Prodh* deficiency does not alter spontaneous excitatory synaptic currents (sEPSCs) of PnC-projecting CeA glutamatergic neurons.**

Representative traces of *in vitro* patch clamp recordings of photo-identified CamKIIα-ChR2 eYFP$^+$ CeA neurons projecting to the PnC, in both WT mice (***left***, black traces) or *Prodh*$^{-/-}$ mice (***right***, red traces). The short (500ms) blue bars above the recording traces represent blue-light pulses used to evoke EPSCs and photo-identify these CamKIIα-ChR2 eYFP$^+$ CeA neurons **B)** Plots showing the cumulative distribution of sEPSC amplitude for WT and *Prodh*$^{-/-}$ mice. N = 7 WT mice; N = 8 *Prodh*$^{-/-}$ mice.

| | WT | *Prodh*$^{-/-}$ |
|---|---|---|
| **Properties of spontaneous EPSCs** | | |
| Events / min | 487.9 ± 170.4 | 689.9 ± 290.9 |
| Amplitude (pA) | 9.7 ± 1.0 | 9.5 ± 1.2 |
| Rise time (ms) | 0.8 ± 0.02 | 0.72 ± 0.05 |
| Decay time (ms) | 4.1 ± 0.89 | 3.12 ± 0.54 |
| Charge (FC) | 20.7 ± 3.4 | 25.4 ± 5.1 |
| N | 13 - 14 | 13 - 14 |
| **Properties of spontaneous action potentials** | | |
| Action potentials / min | 27.8 ± 15.8 | 23.9 ± 12.6 |
| Width at half height (ms) | 1.49 ± 0.09 | 1.58 ± 0.08 |
| **Rheobase (pA)** | **17.8 ± 6.62** | **-6.1 ± 9.3*** |
| N | 12 - 13 | 11 - 13 |
| **Intrinsic properties** | | |
| Capacitance (pF) | 19.0 ± 2.0 | 18.8 ± 2.0 |
| $R_{Series}$ (MΩ) | 16.9 ± 2.5 | 18.5 ± 1.7 |
| $R_{Membrane}$ (GΩ) | 2.87 ± 0.80 | 4.56 ± 1.87 |
| $R_{Input}$ (GΩ) | 539.7 ± 34.0 | 562.22 ± 77.5 |
| N | 12 - 15 | 10 - 15 |

**Supplementary table 1. Intrinsic and synaptic properties of PnC-projecting CeA glutamatergic neurons recorded in WT and *Prodh*$^{-/-}$ mice**

WT N = 7 animals; 2 males, 5 females. *Prodh*$^{-/-}$ N = 8 animals; 4 males, 4 females. *Unpaired t-Test, $p < 0.048$
